# Structural basis of liver de-targeting and neuronal tropism of CNS-targeted AAV capsids

**DOI:** 10.1101/2025.06.02.655683

**Authors:** Tyler J Brittain, Seongmin Jang, Gerard M Coughlin, Bre’Anna H Barcelona, Izabela Giriat, Fiona Ristic, Nathan Appling, Camille PMA Chossis, Timothy F Shay, Viviana Gradinaru

## Abstract

Crossing the blood-brain barrier while minimizing liver transduction is a key challenge in developing safe adeno-associated virus (AAV) vectors for treating brain disorders. In mice, the engineered capsid PHP.eB shows enhanced brain transduction, while the further engineered CAP-B10 is also de-targeted from astrocytes and liver. Here, we solve cryo-EM structures of CAP-B10 and its complex with AAV receptor (AAVR) domain PKD2, at 2.22 and 2.20 Å resolutions, respectively. These structures reveal a structural motif that hinders AAVR binding, which we confirm by measuring affinities. We show that this motif is transferable to other capsids by solving cryo-EM structures of AAV9-X1 and AAV9-X1.1, without and with PKD2, at 3.09, 2.51, and 2.18 Å, respectively. Using this structural information, we designed and validated novel AAV variants with reduced liver and altered brain cell tropism *in vivo*. Overall, our findings demonstrate that rationally modulating AAVR affinity can alter liver targeting and cellular tropism.

## Introduction

Effective gene therapy for neurological diseases requires gene delivery vectors capable of mediating sufficient transgene expression in the central nervous system (CNS) while avoiding immunogenicity^1–3^. Adeno-associated viruses (AAV) have been FDA approved for the treatment of CNS diseases due to their low pathogenicity and stable gene expression across broad cell types including neuronal cells^4–9^. AAV9 has been a particular target of interest in engineering due to its natural ability to penetrate the blood-brain barrier (BBB) after systemic (intravenous) delivery^10,11^. However, AAV9 and other naturally occurring AAVs show poor efficiency in crossing the BBB and transduce many tissues body-wide, most problematically resulting in hepatotoxicity^12–16^. This limits the application of AAV-based gene delivery as a potential therapy for treating brain diseases.

Directed evolution strategies have shown that these properties of AAV capsids are highly modifiable, suggesting that engineered vectors can address these issues^17,18^. High- throughput screening strategies have yielded multiple capsids which have been widely adopted in both the basic sciences and preclinical research. AAV9-derived capsid variant libraries are commonly made by inserting a random 7-mer sequence between residues 588 and 589 of the surface exposed variable region (VR)-VIII^19–21^. *In vivo* selections with this library design yielded the brain-enhanced capsid AAV-PHP.B^20^ and subsequent engineering of flanking amino acids yielded the now-widely-used AAV-PHP.eB (PHP.eB)^22^, which is highly enriched in the CNS and moderately de-targeted from other organs in mice after intravenous administration. Additional engineering of PHP.eB at VR-IV by substitution of residues 452-458 with a randomized 7-mer identified a further improved version, AAV.CAP-B10 (CAP-B10)^23^, which more effectively de-targets from the liver in mice. Furthermore, CAP-B10 shows more neuronal and less astrocytic tropism in mouse CNS, suggesting that VR-IV engineering may also influence brain cell type tropism. A separate VR-VIII library selection found the capsid AAV9-X1, which is highly endothelial specific in the rodent brain^24^. By adopting the same VR-IV substitution of CAP-B10, AAV9-X1 was further improved to AAV9-X1.1, which possesses better production yield and stability, while maintaining a similar specificity for brain endothelial cells and a comparable level of liver transduction.

These variants were optimized in mice, however, and the design of similarly enhanced vectors for clinical application is hindered by our limited understanding of AAV infection mechanisms across organs and species, with only a few naturally-evolved receptor-AAV interactions identified to date. For example, terminal N-linked galactose on the cell surface is well known as a primary attachment factor for AAV9 and its derivatives ^25–27^. Another crucial factor is the AAV receptor (AAVR)^28–30^. AAVR comprises five repeated immunoglobulin (Ig)-like polycystic kidney disease domains (PKD1–5), all sharing a conserved β-barrel structure. Each PKD domain presents distinct residues on its surface, facilitating unique interactions. For instance, among PKD1-5, PKD2 exclusively interacts with AAV9^21^. Since AAVR is expressed on the cell surface across various organs and tissues^31–35^, AAVR was thought to be the primary receptor through which AAVs are internalized. However, recent studies have shown that certain serotypes, such as AAV4 and AAVrh32.33, can bind the cell surface and enter cells without AAVR, but cannot transduce the cell^29^. This suggests that AAVR’s role may be more complicated and potentially extend beyond AAV internalization^29^.

Current engineering efforts aim to guide AAVs away from the liver, primarily focusing on ablating capsid glycan binding, which decreases basal AAV infectivity across organs^26,36,37^. However, some of these modifications also make AAV transduction-deficient, stressing the importance of a trade-off between off-target reduction and on-target efficiency. Recent structural studies from our group and others showed that the moderately liver de-targeted PHP.eB has reduced affinity for PKD2 compared to AAV9^21,38^. Cryo-electron microscopy (cryo-EM) structures of PHP.eB alone^21^ and in complex with PKD2^38^ revealed that PHP.eB gained a new hydrophilic interaction between residues in VR-VIII, which we termed the “lysine lever”, that rigidly orients the loop; we suggest that this causes a steric clash with AAVR-PKD2^21^. This observation suggests that liver transduction may be governed by AAVR-PKD2 affinity.

In this study, we trace the directed-evolution path of AAVs from AAV9 to PHP.eB to CAP-B10, the most liver de-targeted variant in its lineage to date, using structural and biophysical methodologies. Based on this, we present seven cryo-EM structures of CNS-targeting and/or liver de-targeted AAVs, and their AAVR binding affinities. Based on the resulting structural insights, we rationally designed novel AAVs with limited liver transduction. Together, our results reveal the mechanism by which directed evolution has changed receptor affinity and transduction selectivity, which should aid in the rational design of engineered AAVs with enhanced liver de-targeting.

## Results

### The VR-IV modification of CAP-B10 disrupts interaction with AAVR-PKD2

The brain-enhanced capsids PHP.eB and CAP-B10 were sequentially derived from AAV9 along a directed evolution path (**Figure 1a-c**, **Table 1**). This generated CAP-B10 from PHP.eB, which introduced neuronal cellular specificity and enhanced liver de-targeting. The underlying mechanism is unclear, but a clue may be inferred from the structure of PHP.eB in complex with its receptor^38^, showing how a “lysine lever” interaction in the VR-VIII 7-mer introduces a steric clash that partially destabilizes the capsid’s interaction with AAVR-PKD2 (**Figure 1c, Supplementary** Figure 1a)^21^. Extending this observation may help explain CAP- B10’s significantly lower liver infectivity. We hypothesized that the VR-IV 7-mer mutation in CAP-B10 (^452^NGSGQNQ^458^ to ^452^DGAATKN^458^) may further disrupt AAVR-PKD2 binding, and thereby further reduce off-target, especially liver, transduction. To investigate whether the VR-IV mutation in CAP-B10 impacts AAVR affinity, we determined high-resolution cryo-EM structures of three capsids: CAP-B10 (2.22 Å resolution), AAV9-X1 (3.09 Å) and AAV9-X1.1 (2.51 Å). We also solved the structures of CAP-B10 and AAV9-X1.1, which share the same VR-IV sequence motif, in complex with AAVR-PKD2 (2.20 and 2.18 Å, respectively) (**Figure 1d, Supplementary** Figures 2**, 3**). In both structures of receptor-capsid complexes, PKD2 domains are associated with the capsid, binding the plateaus around the 3-fold axis, as seen previously in PKD2- complexed structures of AAV9 and PHP.eB^19,21,38^. PKD2 appears to engage the VR-VIII, VR-IV, and VR-I loops of the capsid as a binding platform (**Figure 1e, Supplementary** Figure 1b). We did not observe a difference in PKD2 position between AAV9-X1.1 and CAP-B10, indicating that the 7-residue difference in VR-VIII does not significantly alter the general binding interface.

**Figure 1:**
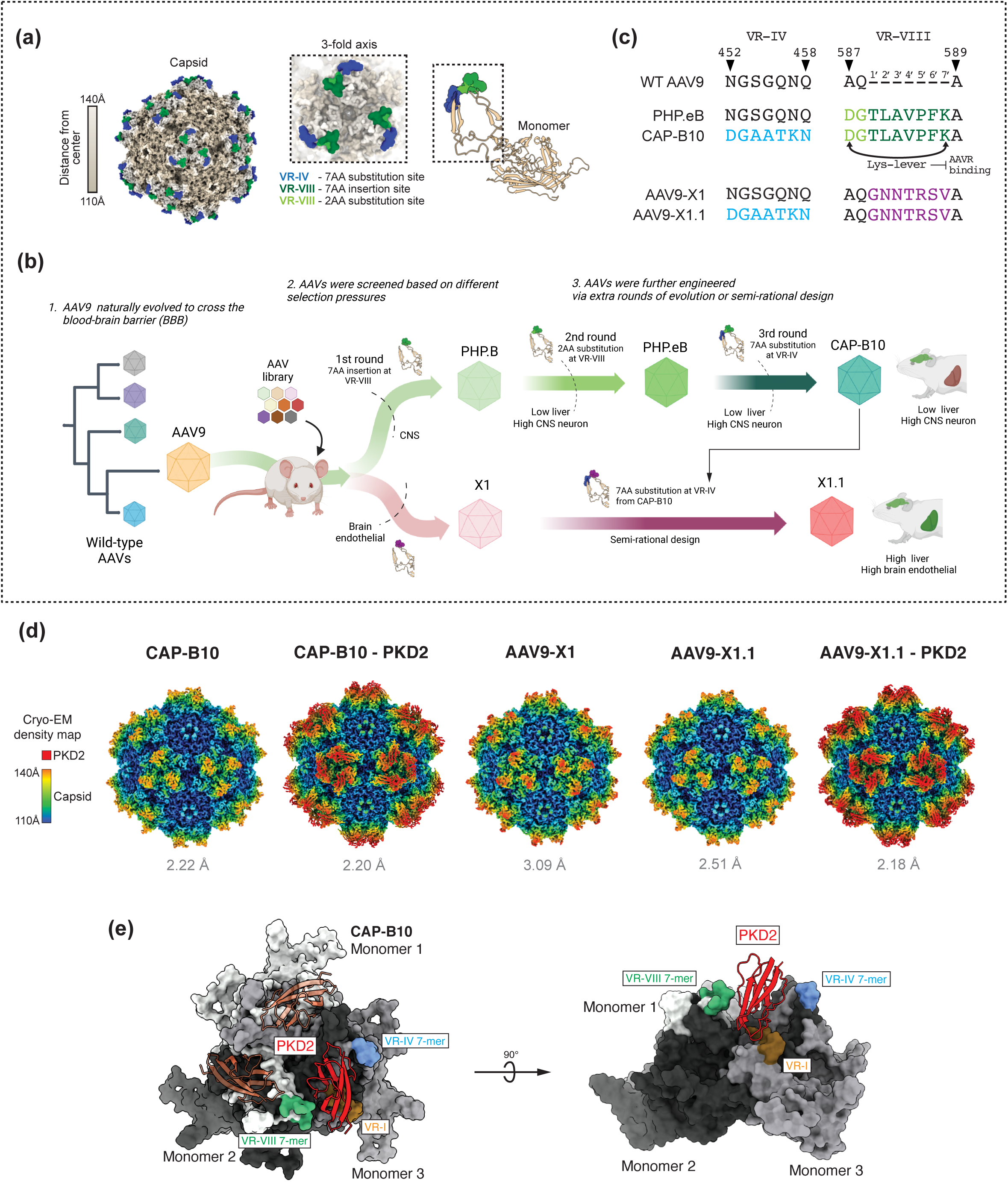
Structure of engineered CNS-enhanced AAVs. **(a)** Location of engineered sites on the AAV capsid shown on structure of PHP.eB (PDB: 7UD4^21^). **(b)** Schematic of directed evolution efforts yielding capsids with high transduction of rodent CNS and low transduction of liver (PHP.B, PHP.eB, and CAP-B10; green arrows), and high transduction of brain endothelial cells and liver (AAV9-X1 and AAV9-X1.1; red arrows). **(c)** Sequence alignment of VR-IV and VR-VIII regions in engineered capsids. The CAP-B10 7-mer substitution is shown in blue, the PHP.eB 7-mer insertion in green, two point mutations in light green, and the AAV9-X1 7-mer insertion in purple. D587 and K7’ in PHP.eB and CAP-B10, form the “lysine-lever” which diminishes AAVR-PKD2 binding. **(d)** Single-particle cryo-EM maps of engineered capsids with and without PKD2, the AAV9 binding domain of AAVR. In the maps, color indicates distance from the center of the capsid. **(e)** Binding interface between PKD2 and 3-fold face of AAV. AAVR-PKD2 is red, VR-I is orange, 7-mer substitution in VR-IV is blue, 7-mer insertion in VR-VIII is green. The individual monomers of the AAV 3-fold face are colored white, light gray, and dark gray.

Since our structures confirmed that the VR-IV loop, mutated in CAP-B10 and AAV9-X1.1, is closely apposed to AAVR-PKD2, we next compared the binding affinity of CAP-B10 for AAVR-PKD2 with that of its ancestral capsids, AAV9 and PHP.eB, measuring equilibrium dissociation constants (K_d_) by biolayer interferometry (BLI) (**Figure 2a, Supplementary Table 2**). AAV9 and derivatives were loaded onto the BLI biosensor and PKD2, serving as the binding analyte, was introduced to the sensor surface at varying concentrations. CAP-B10 interacted with AAVR-PKD2 with a K_d_ of 60.63 µM, indicating a weaker binding affinity than PHP.eB (K_d_: 14.22 µM) (**Figure 2b**) and AAV9 (K_d_: 11.40 µM, consistent with previously reported findings^21^). These results confirm that the 7-residue substitution in the VR-IV region of CAP-B10 disrupts its interaction with AAVR-PKD2.

**Figure 2:**
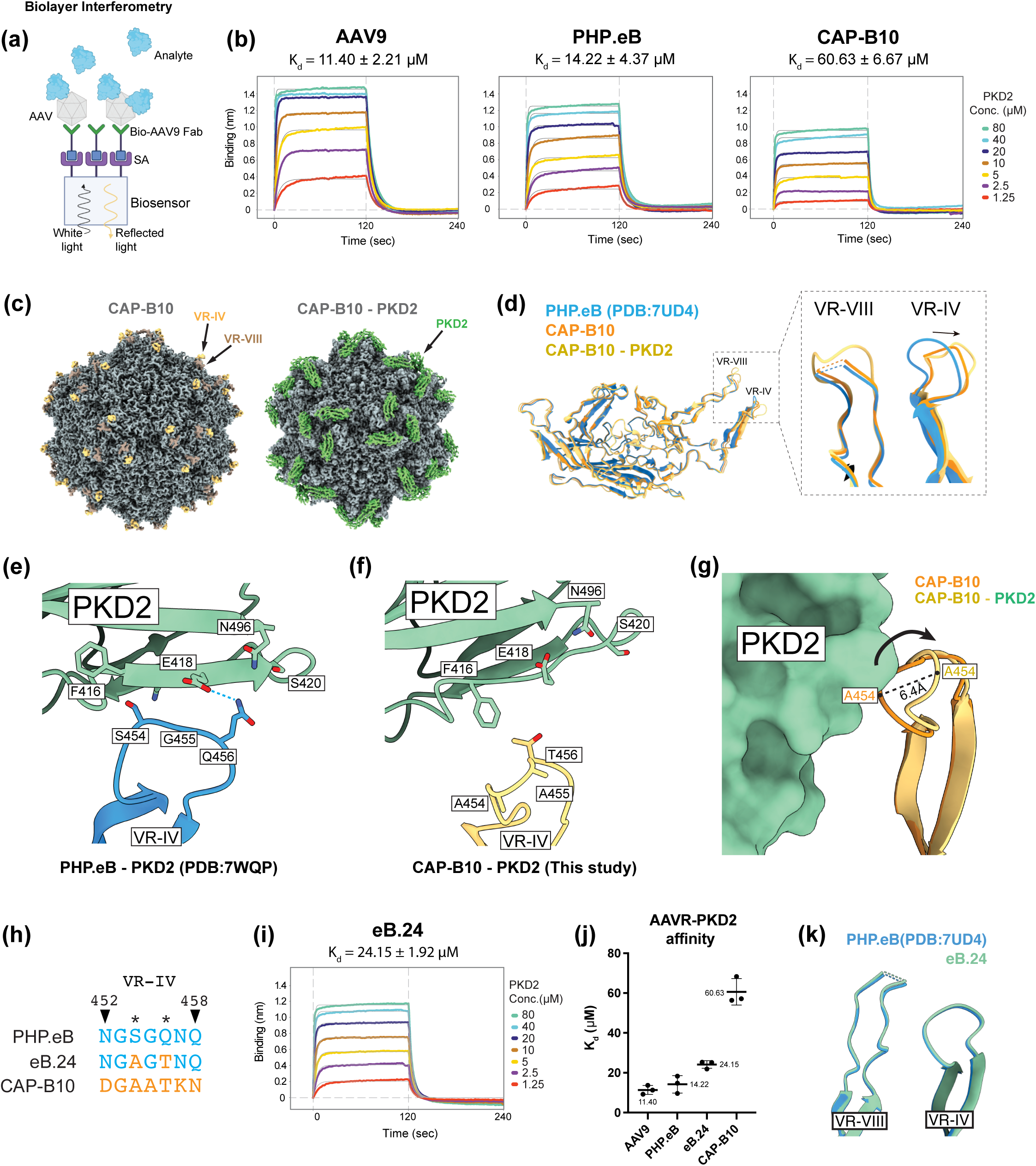
VR-IV modification in CAP-B10 reduces AAVR-PKD2 affinity. **(a)** Schematic of biolayer interferometry (BLI) to measure the binding affinity between AAV capsids and AAVR-PKD2 domain. Biotinylated AAV9 affinity ligand is immobilized on a streptavidin-coated biosensor, AAVs are loaded as the analyte, and binding is quantified by sensorgrams. **(b)** Representative sensorgrams detail the interactions between AAV9, PHP.eB, and CAP-B10 with AAVR-PKD2. Assays were performed three times, with one representative dataset shown. The averaged equilibrium dissociation constant (K_d_) is presented for each capsid. **(c)** Cryo-EM maps of CAP- B10 without and with PKD2. In the uncomplexed map, VR-IV density is shown in orange and VR-VIII density in brown. In the complex map, PKD2 density is shown in green. **(d)** Atomic models of the VP3 subunit of PHP.eB (PDB: 7UD4), CAP-B10 (this study), and CAP-B10 complexed with PKD2 (this study), overlaid to show structural differences and similarities. Inset shows (left) that the structure of VR-VIII in PHP.eB and CAP-B10 is similar, and in the presence of PKD2 additional residues become resolvable, and (right) VR-IV loop structural differences between CAP-B10 (orange) and PHP.eB (blue). Arrow indicates the lower position of VR-IV in CAP-B10 relative to PHP.eB, and the additional conformational change when CAP-B10 is bound to PKD2 (yellow). **(e)** PHP.eB has favorable amino acids at positions 456 and 454 to interact with PKD2. Q456 on PHP.eB can form a hydrogen bond to PKD2 E418 and N496, with PKD2 S420 and PHP.eB S454 aiding in creating a hydrophilic environment. Bulky F416 on PKD2 is positioned away from these hydrophilic residues thereby preventing steric clashes. Dashed sky blue line indicates potential hydrogen bonding **(f)** Our CAP-B10 PKD2 complex structure reveals that two key mutations, S454A and Q456T, play a key role in disrupting the PKD2 interaction. Short T456 prevents hydrophilic interaction with E418, N496, and S420 on PKD2. A454 sterically clashes with F416 on PKD2, causing the VR-IV loop to bend away from PKD2. In addition to these two residues, the generally downwardly bent structure of VR-IV further distances T456 from hydrophilic PKD2 residues. **(g)** Comparison of structures of CAP- B10 (orange) and CAP-B10 complexed with PKD2 (yellow) shows that PKD2 induces a conformational change in the VR-IV of CAP-B10. The greatest shift occurs at A454, which is shifted 6.4Å farther away from PKD2. **(h)** VR-IV sequences of PHP.eB, eB.24, and CAP-B10. The two point mutations in eB.24 are highlighted in orange, S454A and Q456T. **(i)** Representative BLI sensorgram of eB.24 binding of AAVR-PKD2. **(j)** Affinity values of engineered capsids for AAVR-PKD2. Kd values are an average of 3 replicates. **(k)** Cryo-EM- based atomic model of eB.24 overlaid with the structure of PHP.eB (PDB: 7UD4). The two point mutations (S454A, Q456T) in eB.24 do not modify the VR-IV backbone structure of PHP.eB.

### Cryo-EM structure of CAP-B10 complexed with AAVR-PKD2 reveals that hydrogen bonding mediates VR-IV interaction with PKD2

To gain detailed structural insight into the mechanism responsible for the reduced CAP- B10 interaction with PKD2, we delved into our cryo-EM structures of CAP-B10 (**Figure 2c-g**). Electron density was resolved for the side chains of all residues except A454, which is highly solvent exposed, allowing us to fit a model^39,40^(**Supplementary** Figure 2b). We first analyzed structural differences in the VR-IV loop of CAP-B10 (^452^DGAATKN^458^) compared to AAV9 and PHP.eB (^452^NGSGQNQ^458^). Compared to PHP.eB, the VR-IV loop of CAP-B10 is angled away from the capsid surface (**Figure 2d**). A previously-solved structure of the PHP.eB–PKD2 complex showed that the VR-IV loop contributes to the interaction interface (**Figure 2e**). The PKD2 interaction surface is hydrophilic and acidic, and VR-IV accesses this region through hydrophilic residues. For example, Q456 of PHP.eB forms a hydrogen bond with E418 on PKD2 which appears to be a key contributor to the interaction between PHP.eB and PKD2 (**Figure 2e, Supplementary** Figure 4a). Rotamer screening revealed that N496 on PKD2 can potentially form hydrogen bonds as well, although the likelihood is lower than that of PKD2 E418 (**Supplementary** Figure 4b.) Additionally, the bulky F416 on PKD2 is angled away from S454 on PHP.eB, avoiding any steric hindrance and helping S454 and Q456 of PHP.eB form a hydrophilic connection with E418, S420, and N496 of PKD2. In contrast, our CAP-B10–PKD2 complex structure shows that the hydrophilic binding interface is severely disturbed by the Q456T mutation, which abolishes the hydrogen bond with PKD2, and the S454A mutation that replaces a polar serine with a hydrophobic alanine (**Figure 2d, f**). Highlighting the disruption of the hydrophilic interaction is the rotation of PKD2’s F416 into the binding interface and the large ∼6.4 Å conformational shift of the VR-IV loop away from PKD2 (**Figure 2g**).

To verify that the S454A and Q456T mutations in the VR-IV loop are key to reduced PKD2 binding, we created a variant capsid (named eB.24) with only these two residues mutated from the PHP.eB sequence (^452^NG**A**G**T**NQ^458^, bold residues indicate mutations). As predicted, eB.24 showed weaker binding affinity for PKD2 (K_d_ 24.15 µM) by BLI, confirming that these residues significantly influence PKD2 binding (**Figure 2h-j**). To explain the intermediate level of PKD2 binding in eB.24, we solved a cryo-EM structure of eB.24 at a resolution of 2.05 Å (**Figure 2k, Supplementary** Figure 2**, 4c-e**) to compare its structural features with PHP.eB and CAP-B10. This comparison revealed that the backbone structures of VR-IV and VR-VIII in eB.24 are nearly identical to those of PHP.eB (**Figure 2k**), indicating that the two point mutations at residues 454 and 456 are not sufficient to alter the VR-IV backbone conformation and do not affect the structure of VR-VIII either. These findings suggest that the binding differences between eB.24 and PHP.eB stem from the introduced mutations themselves rather than an altered loop conformation, whereas the dramatic difference in PKD2 affinity between PHP.eB and CAP-B10 results from combined alterations in bonding interactions and overall loop conformation.

We next focused on VR-VIII of CAP-B10, which retains the 7-mer insertion and two point mutations (^587^DGTLAVPFK^7’^) from PHP.eB (we denote the amino acids in the CAP-B10 and PHP.eB 7-mer as T1’, L2’, A3’, V4’, P5’, F6’, and K7’). As expected, the VR-VIII of CAP-B10 exhibits a downwardly bent structure similar to that observed in PHP.eB (**Figure 2d, Supplementary** Figure 1). As in PHP.eB, a hydrogen bond formed by D587 and K7’ induces inward tension that bends the 7-mer downward toward the capsid surface. As discussed above, this causes a steric clash with PKD2 by pushing the hydrophobic patch ^2’^LAV^4’^ toward the hydrophilic surface of PKD2, thereby reducing AAVR binding. CAP-B10 therefore incorporates two features, the sterically hindering loop of VR-VIII inherited from PHP.eB and the less hydrophilic loop of VR-IV unique to CAP-B10, that together explain its even weaker affinity for AAVR-PKD2.

Next, we examined whether the VR-IV modification impacts other previously-identified PHP.eB and AAV9 receptors, such as LY6A and D-galactose. We performed pull-down assays, using agarose beads charged with each respective receptor as bait, and analyzed the amount of AAV captured (**Supplementary** Figure 5a). As a control, we first tested the interactions of AAV9, PHP.eB, and CAP-B10 with AAVR-PKD2 to validate the consistency of this method with BLI results (**Supplementary** Figure 5b**, c**). As expected, AAV9 showed the highest PKD2 capture efficiency, while CAP-B10 captured markedly less PKD2, confirming that the pull-down assay results align with BLI results.

Next, we assessed LY6A and D-galactose binding (**Supplementary** Figure 5d-e). Unlike AAVR-PKD2, there was no large difference in receptor binding affinity among PHP.eB, eB.24, and CAP-B10. We observed that CAP-B10 was captured by D-galactose slightly less than PHP.eB or eB.24, but this trend was much less pronounced than the decreasing pattern observed in PKD2 binding from PHP.eB to CAP-B10. This suggests that the CAP-B10 structural motif selectively alters PKD2 affinity while not altering other interactions. It follows then that the tropism differences between PHP.eB and CAP-B10 are primarily driven by the difference in PKD2 affinity, since LY6A and D-galactose interactions remain largely unchanged.

### AAV9-X1.1 VR-VIII counteracts diminished AAVR binding mediated by the CAP-B10 VR-IV motif

After confirming the AAVR hindrance effect of the CAP-B10 7-mer, we investigated whether this property could be transferred to other capsids. To test this, we examined AAV9-X1 and AAV9-X1.1, which selectively target endothelial cells in the rodent brain (**Figure 1**). Unlike CAP-B10, these vectors target the CNS through the LRP6 receptor which, unlike LY6A, is also expressed in the liver^33,34,41–43^, where it contributes to vector transduction^44^. Since liver de- targeting was not selected during engineering, AAV9-X1 exhibits high liver infectivity. AAV9- X1.1 was produced by introducing the CAP-B10 7-mer into VR-IV of AAV9-X1, improving production yield, but not decreasing liver transduction. To test whether the introduced CAP-B10 7-mer weakened PKD2 binding affinity, we performed BLI (**Figure 3a, b**). Surprisingly, AAV9-X1 exhibited a K_d_ of 5.87 µM, stronger than wild-type AAV9 (K_d_ 11.40 µM), indicating that the VR-VIII of AAV9-X1 does not disrupt AAVR-PKD2 binding like that of PHP.eB (K_d_ 14.22 µM). However, the K_d_ of AAV9-X1.1 was 18.85 µM, approximately three-fold weaker than AAV9-X1, suggesting that the diminished AAVR affinity of the CAP-B10 7-mer is transferable. Without the additional destabilization of the interaction from the VR-VIII loop, the affinity of AAV9-X1.1 for PKD2 was comparable to wild-type AAV9, and significantly stronger than CAP-B10 (K_d_ 60.63 µM).

**Figure 3:**
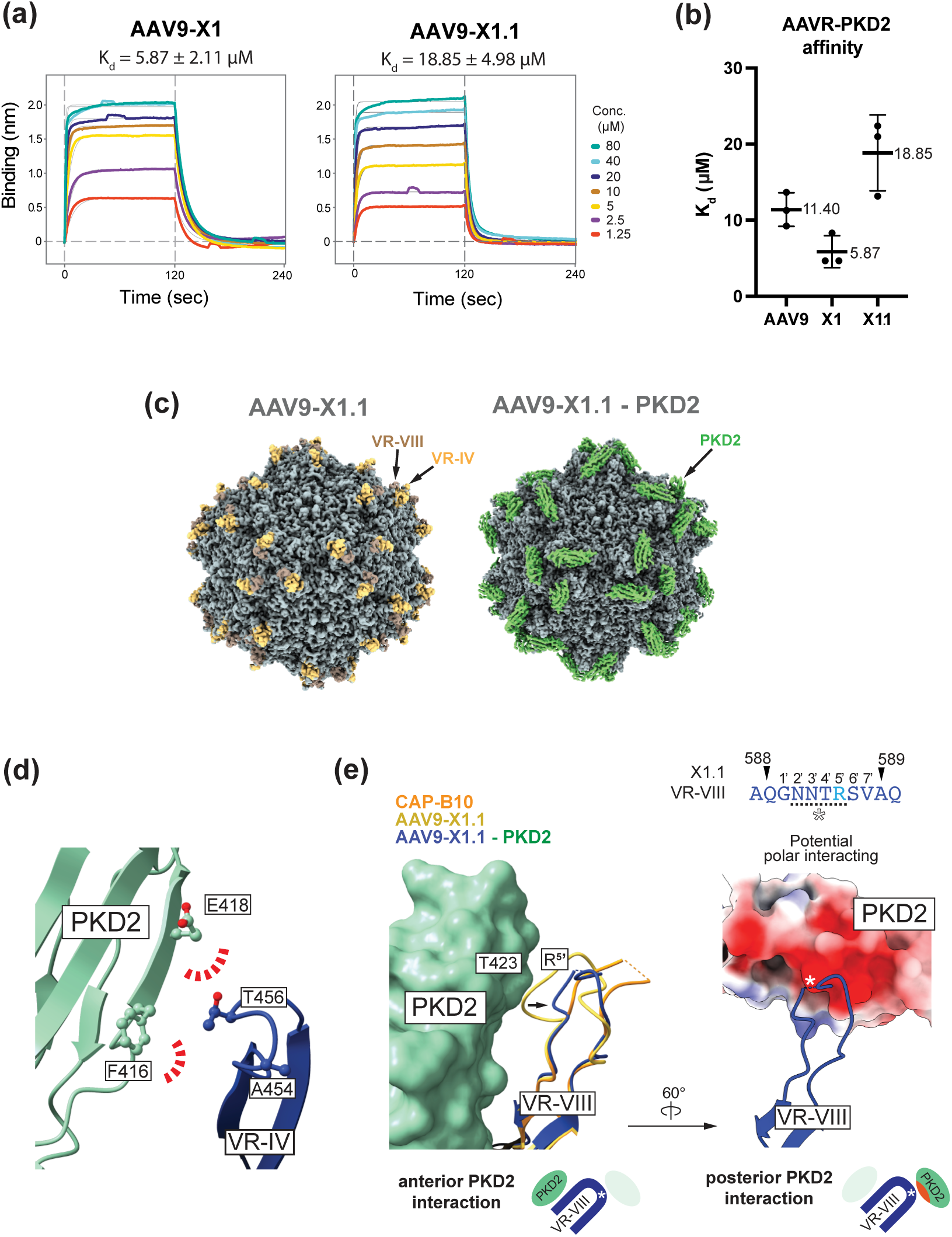
Structural and biophysical characterization of AAV9-X1 and AAV9-X1.1 binding to AAVR- PKD2. **(a)** Representative BLI sensorgrams showing the interaction of AAV9-X1 and AAV9-X1.1 with AAVR-PKD2. Kd values are averages of three replicates. **(b)** Comparison of AAVR-PKD2 binding affinity of AAV9, AAV9-X1, and AAV9-X1.1. AAV9-X1.1 shows weaker AAVR-PKD2 binding affinity compared to AAV9-X1, indicating that the CAP-B10 VR-IV structural motif introduced in AAV9-X1.1 reduces AAVR-PKD2 binding, similarly to CAP-B10. Note that the AAV9 - AAVR-PKD2 affinity shown here is from the same experiment as in Figure 2j. **(c)** Cryo- EM maps of AAV9-X1.1 in its unbound and AAVR-PKD2 bound states. In the uncomplexed structure, the VR-IV is shown in orange and VR-VIII is in brown. In the complex map, PKD2 density is highlighted in green. **(d)** Atomic model of VR-IV in AAV9-X1.1 (blue) complexed with PKD2 (green). As in CAP-B10, T456 loses its hydrogen bond to PKD2 E418, while A454 is pushed away due to a steric clash with PKD2 F416. **(e)** Atomic model of VR-VIII in AAV9-X1.1 (blue) complexed with PKD2 (green surface), overlaid with AAV9-X1.1 (yellow) and CAP-B10 (orange). The schematics below illustrate the interactions between PKD2 and VR-VIII of AAV9- X1.1. *Left:* Interaction of VR-VIII with the anterior PKD2. *Right*: Interaction of VR-VIII with the posterior PKD2. Asterisks denote the four polar-basic amino acids (^2’^NNTR^5’^) at the protruding end of the X1 7-mer that potentially interact with the polar-acidic surface of PKD2.

To further understand how AAVR affinity is maintained in AAV9-X1.1 despite the B10 VR-IV 7-mer, we explored our cryo-EM structures of AAV9-X1.1 with and without AAVR-PKD2, building atomic models of AAV9-X1, AAV9-X1.1, and the AAV9-X1.1–PKD2 complex, at resolutions of 3.09 Å, 2.51 Å, and 2.18 Å, respectively (**Figure 3c-e, Supplementary** Figure 6a-c). AAV9-X1 and X1.1 both contain the VR-VIII 7-mer insertion ^587^AQGNNTRSV^7’^ (we denote the amino acids between 588 and 589 as G1’, N2’, N3’, T4’, R5’, S6’, V7’), differing from PHP.eB and CAP-B10 (^587^DGTLAVPFK^7’^) (**Figure 1b, c**). To analyze structural differences, we superimposed all three structures with CAP-B10 (**Supplementary** Figure 6b**, c**). AAV9-X1 and AAV9-X1.1 exhibit the same VR-VIII geometries as predicted, while AAV9-X1.1 VR-IV adopts the same geometry as in CAP-B10, indicating that the CAP-B10 structural motif is transferable independently of VR-VIII sequence. Analysis of the AAV9-X1.1 VR-IV loop in the PKD2- complexed structure showed that the CAP-B10 7-mer in AAV9-X1.1 also disrupts the AAVR- PKD2 interaction, confirming that AAVR affinity is reduced through a similar molecular mechanism (**Figure 3d, e**). Similar to CAP-B10, we observed a conformational change when binding PKD2 where the VR-IV loop A454 is pushed away from PKD2 roughly 6.4Å (**Figure 2g, 3d**). However, unlike CAP-B10, the VR-VIII of AAV9-X1.1 is not as rigid as the lysine-lever- induced VR-VIII motif (**Figure 1c**), indicated by a conformational change to a lower energy state in AAV9-X1.1 in complex with AAVR-PKD2. An anteriorly-bound AAVR-PKD2 pushes the VR-VIII of AAV9-X1.1 into a position where four flexible groups, ^2’^NNTR^5’^, composed of polar and positive residues, can potentially interact with a negatively charged patch on a posteriorly-bound AAVR-PKD2 (**Figure 3e right**). The greater flexibility of the VR-VIII loop and its ability to interact with the negative patch on a posteriorly-bound AAVR-PKD2 can explain mechanistically why the affinity of AAV9-X1.1 for AAVR is higher than that of CAP-B10. This high AAVR-PKD2 affinity of VR-VIII, along with AAV9-X1.1’s interaction with LRP6^44^, which is expressed in the liver^33,34,41–43^, mitigates the liver de-targeting effect introduced by the CAP-B10 VR-IV motif.

### AAVR-dependent liver de-targeting is independent of brain sequestration

AAVR is an essential host factor that mediates cell entry of most AAV serotypes^28,29^ and the significantly diminished interaction between liver de-targeted capsids and AAVR-PKD2 strengthens our initial hypothesis that reduced liver transduction is a result of weaker binding of AAVR-PKD2. PET imaging showed that up to 35% of systemically-delivered PHP.eB is found in the brain in mice^45^, raising the possibility that liver de-targeting may reflect significant sequestration of AAVs from the bloodstream into the CNS. To test whether AAVR affinity modulation affects liver de-targeting for AAVs being developed and used to study and treat tissues outside the brain^31,32,35,36^, we incorporated the CAP-B10 7-mer into VR-IV of wild type AAV9, creating AAV9-B10. This capsid cannot bind the receptor LY6A^46–49^ and therefore is not expected to accumulate in the brain to a significant level. BLI confirmed that AAV9-B10 has lower affinity (K_d_: 46.90 µM) than AAV9 for AAVR PKD2 (**Figure 4a-c**). We then solved the structure of AAV9-B10 by single-particle cryo-EM at a resolution of 3.05 Å (**Figure 4d, Supplementary** Figure 6d-f). The resulting structure showed that the VR-IV loops of CAP-B10 and AAV9-B10 adopt similar geometries (**Figure 4d**). Additionally, the VR-VIII loop of AAV9- B10 was unaffected, retaining its wild-type AAV9 conformation. These results confirm that the AAVR-PKD2 hindrance effect of VR-IV is highly transferable. To test the relative potency of AAV9 and AAV9-B10 in cells expressing AAVR but not LY6A, we conducted a cell infectivity assay in human embryonic kidney (HEK) 293T cells (**Supplementary** Figure 7). AAV9-B10 showed less infectivity than AAV9, confirming reduced transduction from diminished AAVR affinity.

**Figure 4:**
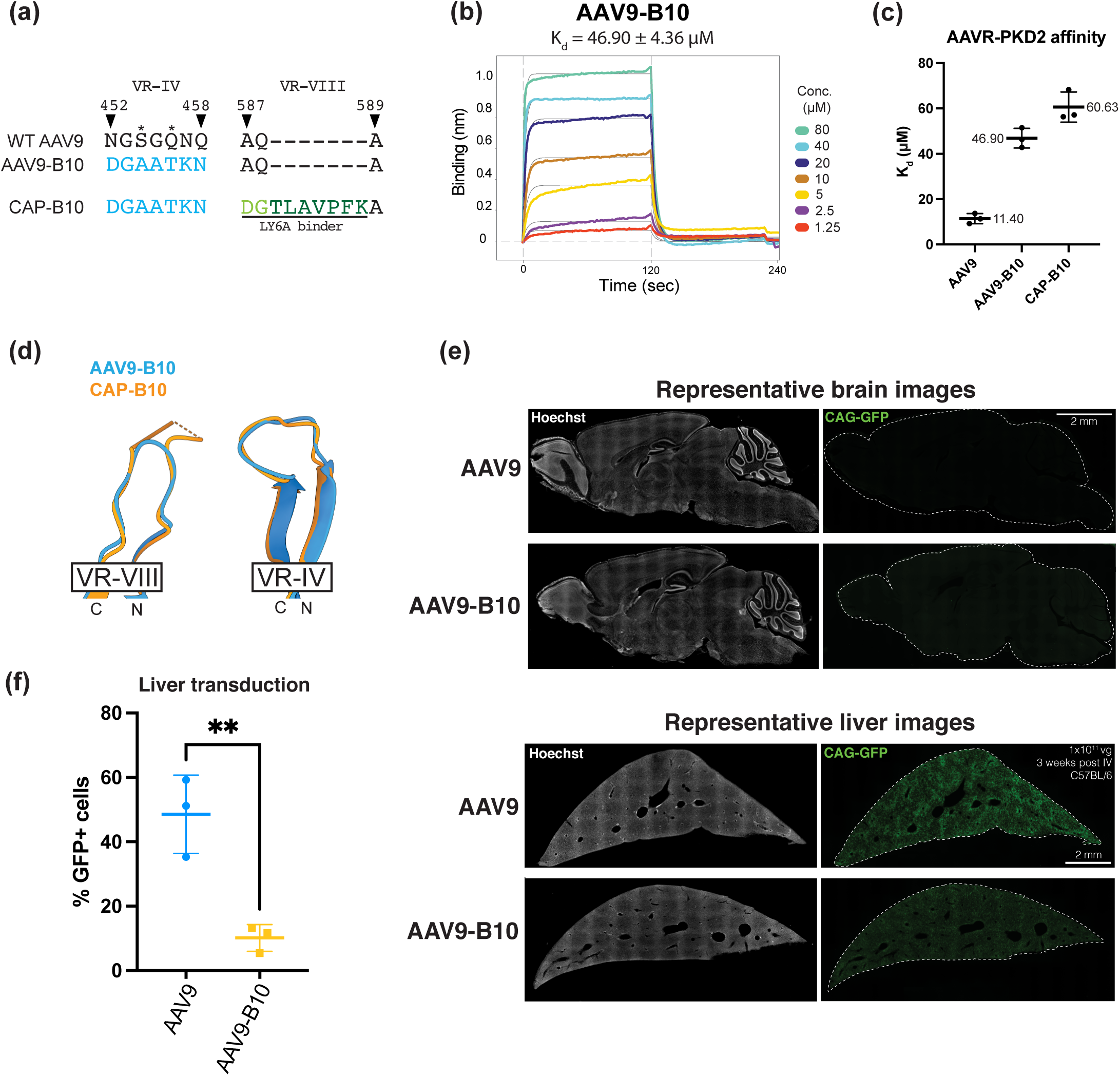
B10 structural motif can reduce liver transduction without brain sequestration. **(a)** Sequence alignment of VR-IV and VR-VIII in AAV9, AAV9-B10, and CAP-B10. AAV9-B10 contains the B10 7-mer substitution in VR-IV but lacks the PHP.eB 7-mer in VR-VIII, preventing interaction with LY6A. **(b)** Representative BLI sensorgram of AAV9-B10 binding to AAVR-PKD2. The averaged K_d_ from triplicate measurements is shown above. **(c)** Comparison of the AAVR- PKD2 affinity values for AAV9, AAV9-B10, and CAP-B10. AAV9 and CAP-B10 data are repeated from Figure 2b and 2j for reference. **(d)** Cryo-EM structure of AAV9-B10, demonstrating that its VR-IV conformation is identical to that of CAP-B10. Additionally, AAV9’s native VR-VIII structure is not altered by the addition of the B10 structural motif at VR-IV. **(e)** Representative images of the brain and liver of mice following systemic delivery of AAV9 or AAV9-B10 packaging eGFP under the control of the CAG promoter. AAVs were retro-orbitally administered to the mice at a dose of 1×10^11^ vg per animal (n = 6 animals per capsid). eGFP expression was analyzed 3 weeks post-injection. Scale bars represent 2 mm. **(f)** Percentage of eGFP-expressing cells in the liver of mice receiving AAV9 or AAV9-B10. AAV9-B10 has markedly decreased liver transduction compared to AAV9 (**p < 0.01). Statistical significance was determined using a two-tailed unpaired t-test.

To test this effect *in vivo*, AAV9 and AAV9-B10 packaging the reporter gene GFP were retro-orbitally injected into wild-type C57BL/6J mice (**Figure 4e**). After 3 weeks, brain and liver transduction levels were assessed. Transgene expression levels were quantified by counting GFP positive cells, normalized to total cell counts determined by Hoechst staining. As expected, liver transduction of AAV9-B10 was markedly decreased compared to AAV9 (**Figure 4f**). Additionally, the GFP levels in the brain were comparable, and low, in both. These results suggest that the CAP-B10 7-mer substitution decreases transduction by reducing AAVR-PKD2 affinity in a manner that is independent of brain sequestration.

### AAVR affinity of brain-enhanced AAVs governs de-targeting from the liver and CNS astrocytes

CAP-B10 is notable both for its liver de-targeting and for its enhanced specificity for neurons in the CNS^23^. To determine if both features arise from CAP-B10’s VR-IV 7-mer-mediated loss of affinity for AAVR, we next tested whether an intermediate AAVR affinity would yield an intermediate vector tropism. To test this hypothesis, we assessed three AAV9 derivatives with varying PKD2 affinities – PHP.eB (K_d_: 14.22 µM), CAP-B10 (K_d_: 60.63 µM), and eB.24 (K_d_: 24.15 µM) – *in vitro* and *in vivo*. As before, we first performed an *in vitro* infectivity assay in HEK293T cells (**Supplementary** Figure 7a**, c-e**). As predicted, eB.24 displayed an intermediate level of infectivity, while PHP.eB and CAP-B10 showed the highest and lowest infectivity, respectively (**Supplementary** Figure 7c**, d**). To see if the presence of an alternate receptor had any effect on transduction, we transfected the LY6A of C57BL/6J mice into HEK293T cells. Interestingly, all LY6A-dependent capsids demonstrated high infectivity in LY6A-expressing cells, with no significant differences between them, despite the substantial differences in AAVR PKD2 affinity among PHP.eB, eB.24, and CAP-B10 (**Supplementary** Figure 7a**, e**). Taken together with the LY6A pulldown experiment (**Supplementary** Figure 5d) these results suggest that reduced AAVR-PKD2 affinity, which reduces infection of the liver, can be compensated for by strong alternative receptors in the brain, such as LY6A.

To test transduction *in vivo*, AAVs were retro-orbitally injected into wild-type C57BL/6J mice, with each capsid packaging GFP as a reporter gene. Liver transduction levels were measured by quantifying cells with GFP fluorescence, normalized to total cell count from Hoechst (**Figure 5a, b**). As expected, PHP.eB showed the highest level of liver transduction, and CAP-B10 the lowest. eB.24, which has less PKD2 affinity than PHP.eB, exhibited significantly lower liver transduction compared to PHP.eB. These results indicate that, indeed, attenuating the AAVR affinity of engineered AAVs can de-target capsids from the liver. We also measured GFP fluorescence in brain tissue to investigate the correlation between brain transduction efficiency and AAVR-PKD2 affinity. Consistent with previous reports^23^, qualitative analysis of brain slices revealed that CAP-B10 exhibited greater neuronal tropism compared to PHP.eB, which transduced both neuronal and astrocytic cells (**Figure 5a**). To further assess the brain cell-type specificity of these viruses, we performed immunohistochemistry using Nissl and S100B, markers for neuronal and astrocytic cells respectively (**Figure 5c, d**). PHP.eB and CAP- B10 achieved similar levels of neuronal transduction, with CAP-B10 exhibiting significantly lower astrocytic transduction compared to PHP.eB, reproducing previous results ^23^. Interestingly, eB.24, which has intermediate AAVR affinity, exhibited neuronal transduction levels comparable to those of PHP.eB and CAP-B10, but exhibited intermediate astrocytic specificity. This observation suggests that AAVR affinity not only modifies liver targeting of AAVs but can also selectively alter cell-tropism in brain-targeted engineered AAVs.

**Figure 5:**
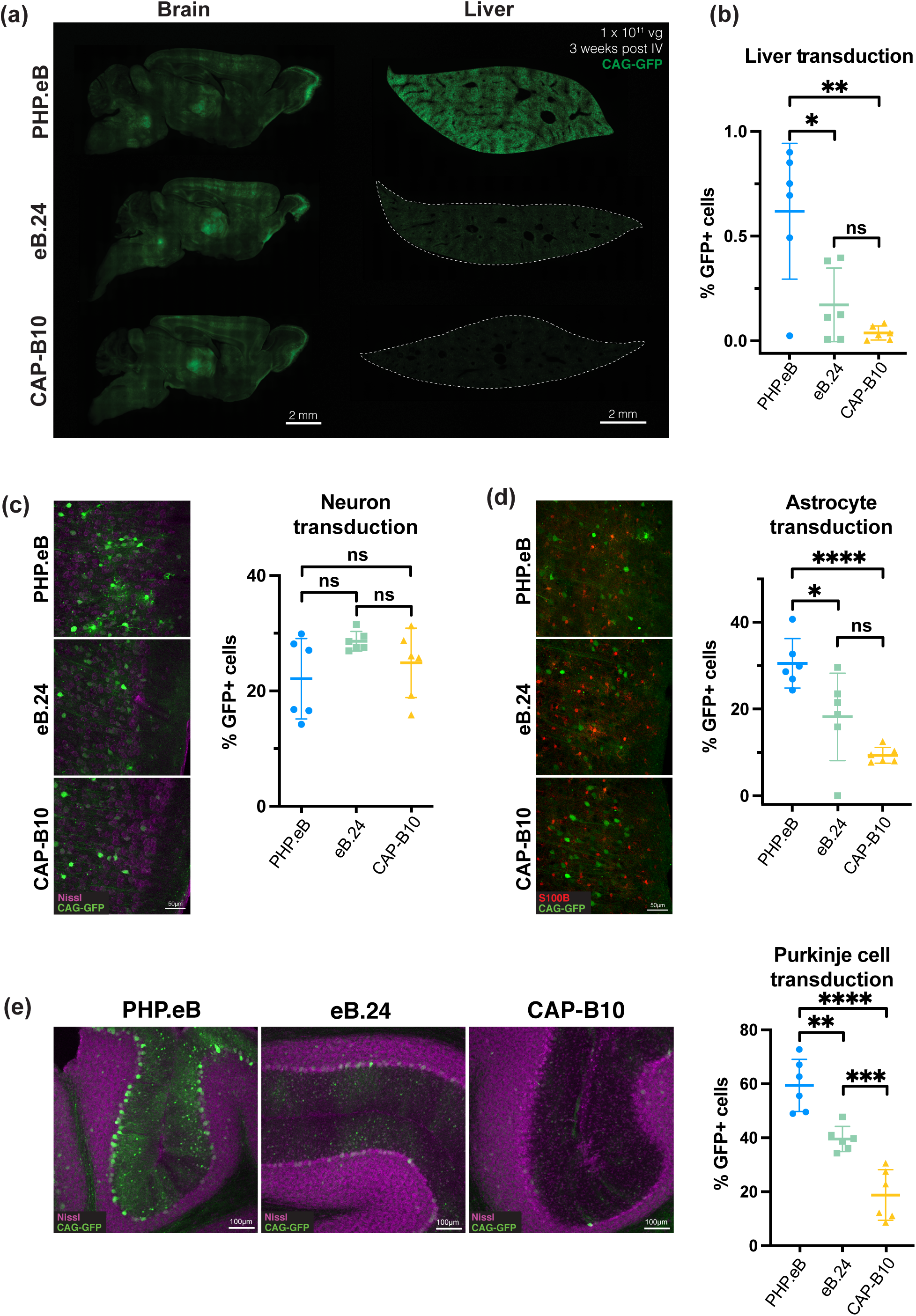
AAVR-PKD2 affinity modulates liver and brain tropism. **(a)** Representative fluorescence images showing eGFP expression in the brain and liver following retro-orbital delivery of PHP.eB, eB.24, or CAP-B10. AAVs packaging the fluorescent reporter eGFP under the ubiquitous CAG promoter were administered retro-orbitally at a dose of 1×10^11^ vg per mouse (n = 6 animals per capsid). eGFP expression was analyzed 3 weeks post-injection. The fluorescent reporter was used to quantify varying levels of brain and liver transduction. Scale bars represent 2 mm. **(b)** Quantification of eGFP-positive cells in the liver shows a correlation with the binding affinity for AAVR-PKD2. These findings demonstrate that liver tropism can be modulated by tuning AAVR-PKD2 affinity. **(c)** Representative images of eGFP-expressing cells co-stained with Nissl, a marker for neurons, to assess neuronal transduction in the cortex. The percentage of GFP+ neurons is consistent between PHP.eB, eB.24, and CAP-B10, suggesting that AAVR affinity does not significantly alter neuronal transduction levels. Scale bar represents 50 µm. **(d)** Percentage of eGFP-expressing cells co-localizing with S100B, indicating astrocyte transduction efficiency. Similar to the liver, astrocytic tropism decreases as AAVR-PKD2 affinity decreases, highlighting the role of AAVR binding in regulating astrocyte transduction. **(e)** *Left:* representative images of eGFP expression in the cerebellum delivered by PHP.eB, eB.24, or CAP-B10 where eGFP is in green and Nissl-stained neurons are in magenta. *Right:* percentage of eGFP-positive Purkinje cells in the cerebellum. As AAVR-PKD2 affinity decreases the transduction level of Purkinje cells decreases. Statistical significance was determined using a two-tailed unpaired t-test. *p < 0.05, **p < 0.01, and ***p < 0.001.

When CAP-B10 was identified, another unique feature was observed: a large reduction in transduction of cerebellar Purkinje cells^23^. To see how well eB.24 emulates this behavior of CAP-B10, we quantified Purkinje cell transduction for PHP.eB, eB.24, and CAP-B10 (**Figure 5e**). As expected, CAP-B10 had much lower Purkinje cell transduction compared to PHP.eB. Interestingly, eB.24 exhibited an intermediate level of transduction between PHP.eB and CAP- B10. The decrease in cerebellar Purkinje cell transduction seen with eB.24 demonstrates that the change in tropism from PHP.eB to CAP-B10 is driven by the decrease in AAVR-PKD2 affinity.

As demonstrated by our cryo-EM structure of eB.24 (**Figure 2k, Supplementary** Figure 4c-e), the difference in PKD2 binding between PHP.eB and eB.24 stems from the two mutations themselves, which do not modify the overall loop conformation. Our *in vivo* transduction assays further imply that these two mutations alone are sufficient to significantly redirect AAV away from the liver, brain astrocytes, and Purkinje cells in the cerebellum. CAP-B10 exhibits a further reduction in AAVR-PKD2 affinity, likely due to an additional structural rearrangement of VR-IV, which further enhances de-targeting of liver, brain astrocytes, and cerebellar Purkinje cells. These results suggest that the effects of AAVR affinity on liver and brain cell-type tropism operate along a gradient, supporting the idea that further engineering may fine-tune tropism and underscoring the potential for optimizing AAV vectors for liver de-targeting and brain cell-type specificity by modulating AAVR affinity (**Figure 6**, **Table 1**).

**Figure 6:**
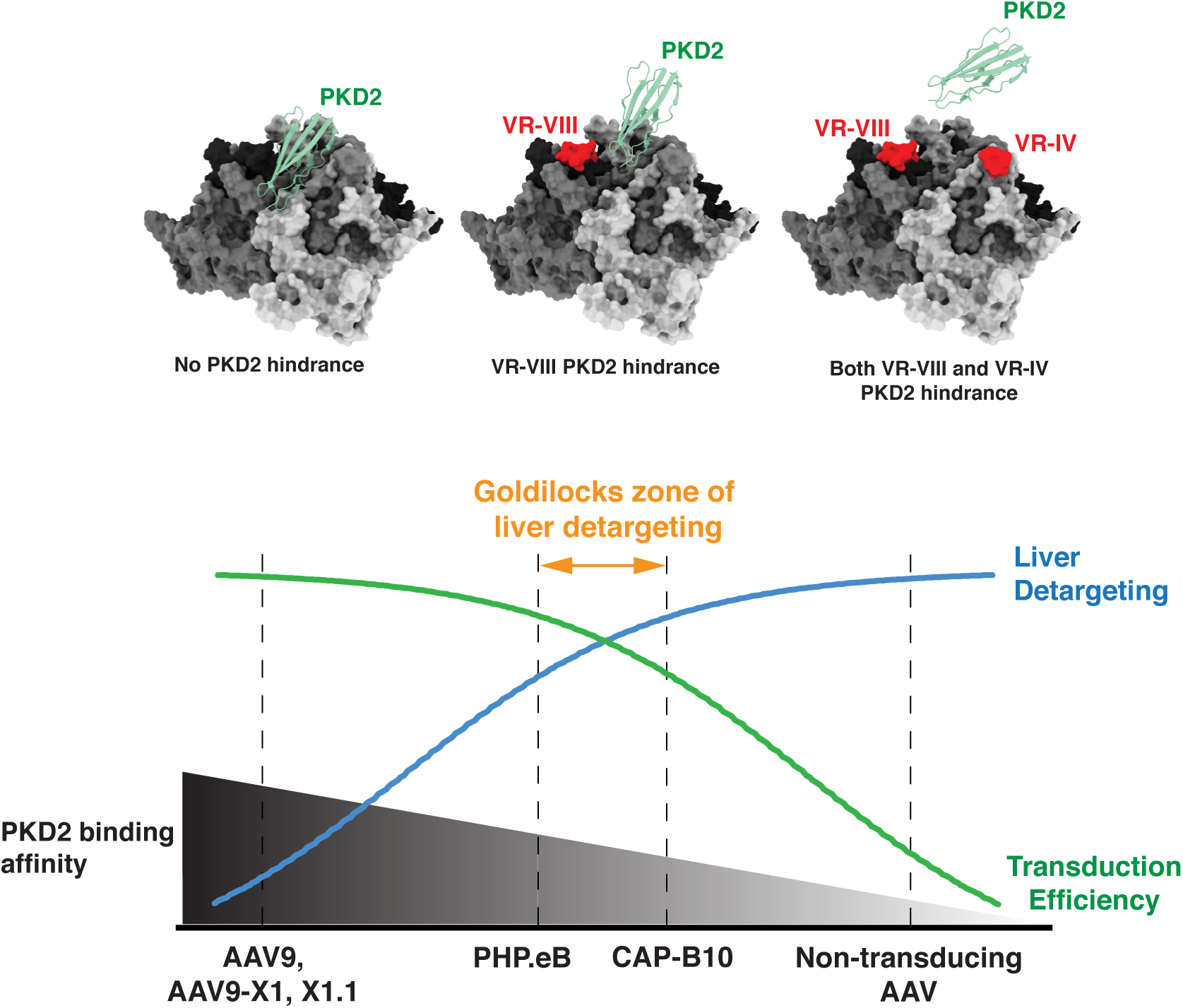
Engineering liver de-targeted AAVs requires a balanced approach. While higher AAVR- PKD2 affinity generally enhances transduction across various organs, it also increases liver transduction. Conversely, AAVs that completely lose AAVR-PKD2 binding fail to transduce any cells. Additionally, even with reduced AAVR-PKD2 affinity, an AAV may still transduce the liver if it engages an alternative liver receptor, such as LRP6 for AAV9-X1.1. Therefore, designing liver de-targeted AAVs requires optimizing AAVR-PKD2 affinity in the context of target-specific receptor interactions.

## Discussion

Directed evolution of AAV capsids has shown that vector transduction patterns across tissues and cell types can be dramatically altered in ways that are potentially advantageous. After intravenous administration, PHP.eB demonstrates broadly potent brain transduction; the more recently identified CAP-B10 focuses that transduction to neurons of the CNS, largely de- targeting the liver (a source of serious adverse events in patients)^50–53^. Understanding the mechanisms behind these capsids’ potentially advantageous transduction properties may accelerate the creation of vectors better suited to human applications. While investigation of broadly potent CNS vectors, like PHP.eB, has yielded a host of new BBB transcytosis receptors, including a subset conserved in humans^44,46,54,55^, mechanisms of neuron specificity and liver de- targeting remain opaque.

Here, based on detailed structural comparisons, binding characterizations, and *in vivo* observations, we reveal that CAP-B10 achieves liver de-targeting through reduced affinity for AAVR. Our analyses of previously reported (AAV9, CAP-B10, X1, X1.1) and newly developed (AAV9-B10, PHP.eB.24) capsids demonstrate that AAVR affinity operates as a dial governing both liver transduction and degree of neuronal specificity (**Figure 6**). Our high-resolution structures of AAVR interactions with engineered AAV capsids also show that fine-tuning of receptor binding is achievable if we understand molecular and structural mechanisms.

There have been efforts to engineer “canvas capsids^36^” with ablated basal infectivity as a starting point for the creation of vectors that might selectively target and/or de-target specific tissues ^26,37,56,57^. The CAP-B10 loop substitution in VR-IV offers a potential strategy for generating such capsids by reducing AAVR-PKD2 affinity^58^. While CAP-B10’s full behavior appears to be explained by the gain of a LY6A interaction and a weakened AAVR interaction, AAVR knockout severely ablates AAV infectivity, reducing it to near zero, and this effect is not rescued by LY6A alone^48^. This explains why CAP-B10 has to maintain a low-but-not-zero AAVR affinity: to reduce off-target tissue infectivity while still being sufficient for AAVR-mediated transduction in some cell types. This highlights the importance of considering receptor binding not as an on/off switch but rather as a control dial, which can and should be tuned during AAV engineering. A recent study showed that AAVR might function as a trafficking factor^29^, implying that a modest affinity which is enough for AAV to remain in proximity to AAVR might be sufficient for AAVR-mediated transduction. However, detailed mechanisms of AAVR and LY6A function during cell infection remain lacking; designing an optimal receptor affinity balance will likely require a deeper understanding of the entire process from AAV uptake and trafficking to eventual nuclear import.

In our previous cryo-EM study^21^, we demonstrated that AAVR affinity can be controlled by altering the insertion length and geometry at VR-VIII. The present work expands this understanding of the AAVR interface to VR-IV, enabling rational affinity adjustments^21^. Through our cryo-EM structures, we find that Q456 of the wild-type AAV9 VR-IV loop is positioned to interact with E418 from AAVR-PKD2 via hydrogen bonding. In CAP-B10, substitution to the more compact T456, along with the overall loop shape bending away from PKD2, disrupts hydrogen bonding, leading to a significant loss of AAVR-PKD2 affinity. Another substituted residue, A454, is pushed away from PKD2 by 6.4Å in the complexed structure, suggesting a steric clash between the hydrophobic alanine and the PKD2 surface. We engineered a new capsid, eB.24, built on the PHP.eB backbone with two point mutations, S454A and Q456T, in the VR-IV loop, which exhibits an AAVR affinity intermediate between those of PHP.eB and CAP-B10. The cryo-EM structure of eB.24 showed that the overall loop shape of AAV9 VR-IV was not altered by the two mutations, confirming that CAP-B10’s affinity change results from both loss of bonding interactions and the introduction of a steric clash.

As expected, eB.24, which has a level of AAVR affinity intermediate between PHP.eB and CAP-B10, had a liver transduction level similarly between PHP.eB and CAP-B10. Additionally, we observed that eB.24 has altered Purkinje cell and astrocyte transduction, suggesting that modulating AAVR affinity can alter cell-type tropism within the brain. The decreased Purkinje cell tropism may result from a different amount of AAVR expression in that cell-type; further research is needed. These observations may explain a recent surprising result from pigtail macaques, where PHP.eB, whose BBB receptor LY6A is not present in non-human primates, exhibited more neuronal tropism than AAV9 following intracerebroventricular administration^59^.

Importantly, the effects of AAVR affinity modulation on liver transduction are not limited to brain-targeting AAVs. AAV9-B10, with its unaltered AAV9 VR-VIII and substituted CAP-B10 VR-IV loop, exhibited significantly lower AAVR affinity and liver transduction in mice than AAV9, and no increase in CNS transduction^20,23^. Thus, this mechanism of liver de-targeting should be broadly applicable to engineered capsids across tissues of interest.

We also tested the binding of LY6A and D-galactose to PHP.eB, eB.24, and CAP-B10 to assess whether the VR-IV modification affects non-AAVR receptors. Overall, the interaction levels remained largely unchanged, except for a slight reduction in D-galactose binding observed with CAP-B10 compared to PHP.eB and eB.24. While VR-IV modification is quite distant from the well-established D-galactose binding site on the 2-fold wall of the AAV surface ^25,26,60^, it is possible that the more polar residues in the protruding VR-IV region of PHP.eB may facilitate a non-specific polar interaction with D-galactose that is lost in CAP-B10. However, the monosaccharide D-galactose beads we used to assay binding do not fully mimic the glycoprotein environment of D-galactose on the cell surface, so further investigation is needed to determine how binding is affected *in vivo*.

Our analysis of a separate family of capsids, AAV9-X1 and AAV9-X1.1, shows that the effect of diminished AAVR affinity can be masked by the contributions of other receptors. AAV9- X1.1 exhibits lower AAVR affinity than AAV9-X1, indicating that the B10 loop hinders AAVR binding in multiple capsid variants. However, AAV9-X1.1’s affinity was higher than CAP-B10’s. This is likely because, unlike for CAP-B10, no structural motifs in VR-VIII appear to disrupt AAVR binding and the VR-VIII of X1/X1.1 may additionally participate in engaging AAVR. AAV9- X1 also uses a different receptor, LRP6, which is enriched both in the brain and liver. Knockout of LRP6 in the liver revealed that this receptor markedly boosts AAV9-X1.1 liver transduction^44^. These observations suggest that the effect of B10’s VR-IV structural motif on AAVR binding, and thus liver de-targeting, is counteracted by LRP6-mediated liver transduction.

Our study presents the first comprehensive structural study of the molecular mechanism by which brain-enhanced AAVs achieve liver de-targeting, providing key insights into their tropism and therapeutic potential. By solving seven cryo-EM structures—five distinct AAVs and two AAV–AAVR-PKD2 complex structures—we reveal the molecular mechanism underlying this process. Our results reveal that CAP-B10 achieves improved liver de-targeting by reducing AAVR affinity through disrupting hydrogen bonds between the VR-IV loop and AAVR-PKD2. *In vivo* testing confirmed that AAVR affinity correlates with liver transduction levels in both CNS- targeting and non-CNS-targeting capsids, and introducing strategic mutations can tune the AAVR affinity to achieve the desired transduction level. Combining VR-IV and VR-VIII modifications could enable the creation of blank canvas capsids where most tissues are de- targeted by tuning AAVR affinity and desired tissues are specifically targeted by engineering an interaction with another receptor. However, optimizing this strategy requires careful selection of new receptors, such as LY6A, which is not enriched in the liver unlike LRP6. While this study focused on rodent CNS-targeting capsids, the mechanistic insights gained should be broadly applicable for future capsid engineering across tissues and species, including humans.

## Methods

### Animals

All mouse experiments were approved by the California Institute of Technology Institutional Animal Care and Use Committee (IACUC). Adult (6-8 weeks old) C57BL/6J WT male mice were purchased from the Jackson Laboratory (000664).

### Viral production

AAVs were produced as previously described^61^. Briefly, HEK293T cells were cultured in 150-mm dishes at 37 °C with 5% CO_2_ until they reached 80%–90% confluency. HEK293T cells were triple-transfected with pHelper, capsid (AAV9, PHP.eB, CAP-B10, eB.24, AAV9-B10, AAV9-X1, AAV9-X1.1), and genome plasmid (eGFP reporter gene controlled by SCP1 promoter for AAV9-X1.1, and eGFP reporter gene controlled by CAG promoter for other else), using polyethylenimine (PEI Max®, Polysciences) solution. Media was collected 72 hr after transfection, and media and cells were collected at 120 hr. AAVs were purified from cells and media through an iodixanol gradient. Purified AAVs were titered using droplet digital PCR (ddPCR, Bio-Rad).

### Protein preparation

The PKD2 domain of AAVR (AA 401–498) was expressed in BL21 (DE3)-RIPL *E. coli* as a fusion protein carrying Myc and 6×His tags. Following cell lysis, insoluble material was removed by centrifugation at 20,000 ×g for 1 hour, and the resulting supernatant was loaded onto an Ni-NTA affinity purification column (Qiagen). The column was pre-equilibrated with 50 mM Tris-HCl (pH 8.0), 100 mM NaCl, and 20 mM imidazole, then washed sequentially with 20 mM Tris-HCl (pH 8.0), 20 mM imidazole, and NaCl gradients (500, 1,000, 500, and 150 mM) to remove non-specifically bound proteins. Elution was performed using 20 mM Tris-HCl (pH 8.0) and 100 mM NaCl, applying an imidazole step gradient (50, 100, 150, and 250 mM), with the 250 mM imidazole fractions collected. PKD2 monomer was separated using Superdex 200 size-exclusion chromatography column (Cytiva) for further experimentation.

For LY6A (AA 1–109) derived from C57BL/6 mice, a construct carrying Fc (human IgG1), Myc, and 6×His tags was transiently transfected into HEK293T cells at 80%–90% confluency using PEI. Conditioned media containing secreted His-Fc-LY6A were harvested 120 hours post-transfection, filtered through a 0.22-µm PES vacuum filter (Sigma-Millipore), and clarified. Ni-NTA resin (Qiagen) was used to capture His-Fc-LY6A, which was subsequently eluted with 100 mM NaCl, 50 mM Tris-HCl (pH 8.0), and 150 mM imidazole. The eluted protein was concentrated using a 10K Amicon concentrator (Sigma-Millipore) before downstream applications.

### Cryo-EM sample preparation and collection

3.5 µL of concentrated AAV (1×10^12^ vg/mL - 1×10^13^ vg/mL) was pipetted onto Quantifoil R 1.2/1.3 300 mesh Cu grids that had been glow discharged (Pelco EasiGlow, 10 mA, 1 min). Grids were then plunge-frozen using a Mark IV Vitrobot (FEI, now Thermo Fisher) (23 °C, 100% humidity, blot force 3, blot time 5 s). Grids were clipped and loaded onto either a 300 kV Titan Krios microscope (Thermo Fisher) equipped with a K3 6k x 4k direct electron detector (Gatan) or a 200 kV Talos Arctica microscope (Thermo Fisher) equipped with a K3 6k x 4k direct electron detector (Gatan). Micrographs were collected using SerialEM software^62^ with pixel sizes of 0.42 Å or 0.85 Å (Arctica and Krios, respectively) and a defocus range of 1.5-4 μm.

### Cryo-EM processing

Processing steps are outlined in **Supplementary** Figure 3. In brief, raw movies were binned by 2 and gain and motion corrected in CryoSPARC(v4.1)^63^. CTF estimation was performed in CryoSPARC. Initial particle picks were generated with blob picker using a maximum diameter of 300 Å and a minimum diameter of 240 Å. Particles were inspected & filtered to eliminate obvious bad picks. The remaining picks were then extracted from micrographs with 2x binning. 2D classification with 100 classes was then used to exclude remaining junk particles. Ab-initio with one class was then run with C1 symmetry. Afterwards, a homogeneous refinement enforcing I1 symmetry was run, aligning most particles at a lower resolution. These particles were then re-extracted unbinned and used for another one class ab-initio with C1 symmetry. The results were fed into a homogeneous refinement with global CTF correction and enforced I1 symmetry. Finally, a non-uniform refinement was run and this map was used for post processing and subsequent model building in Phenix and COOT^39,40,64^.

A model of the VP1 capsid protein was built using AAV9 as a template (PDB: 3UX1^19^). This was fitted into the I1 capsid map using ChimeraX^65^. This was then fed into COOT^64^ and 7- mer insertions and/or substitutions were added or edited and manually fit into the density. The model and map were then iteratively refined in Phenix and manually tuned in COOT for several iterations before a final model was determined (Supplementary Table 1.). Analysis of models was done in ChimeraX.

### Biolayer interferometry

The equilibrium dissociation constant (K_d_) values were determined using Blitz (formerly ForteBio, now Sartorius). A total of 4 μL of 0.2 mg/mL CaptureSelect™ biotin-conjugated AAV9 nanobody (Thermo Fisher^66^) was loaded onto an Octet® SAX Streptavidin-coated biosensor (Sartorius) for 120 s. The biosensor was then incubated in 400 μL of running buffer (Dulbecco’s phosphate-buffered saline [DPBS, Gibco™], 1% BSA, 0.02% Tween-20) to wash out residual nanobody. Next, 4 μL of AAV at 5×10¹² vg/ml was loaded onto the biosensor to form an AAV biolayer. After a 30 s washing step in 400 μL of running buffer, the biosensor was incubated with 4 μL of PKD2 for 120 s (association), followed by incubation in 400 μ: of running buffer for 120 s (dissociation). After each cycle, the biosensor surface was regenerated by incubating it in 400 μL of 10 mM glycine pH 2.0 to remove bound AAV, and the AAV layer was reloaded at the beginning of each new cycle as described above. Sensorgrams were collected at least 6 times in the range of PKD2 concentration between 1.25 - 80 μM. Sensorgrams were analyzed using BLItz Pro software (formerly ForteBio, now Sartorius). K_d_ was measured from 3 experiments and averaged.

### Pull-down assays

Pull-down assays were performed as previously described^21,44^. Briefly, for protein receptor pull-down assay, prey AAVs (0.05 pmol) were mixed with 6xHis-tagged bait, either purified AAVR-PKD2 (100-200 pmol) or His-Fc-LY6A (30–60 pmol) and 15 μL Ni-NTA resin in a DPBS buffer with 20 mM imidazole for 1 hr at 4 °C in a rotary shaker in agitation mode. The mixture was loaded onto a spin column and washed twice with 100 μL (10 column volumes total) DPBS buffer with 20 mM imidazole to remove unbound AAVs. Prey AAVs interacting with the 6xHis-tagged bait were eluted with 60 μL DPBS buffer containing 150 mM imidazole. The resulting eluate was electrophoresed by SDS-PAGE and analyzed by western blotting with anti- VP1/VP2/VP3 (Arp, cat# 03-61058) and anti-6xHis (Abcam, ab18184) antibodies. Western blot images were analyzed using Image Lab (Bio-Rad).

For the D-galactose pull-down assay, D-galactose agarose resin (Thermo Fisher, cat# 20372) was prepared by mixing with Ni-NTA resin at ratios of 1:1, 1:2, and 0. Prey AAVs in DPBS were then mixed with D-galactose resin for 1 hr at 4 °C on a rotary shaker in agitation mode. Resin was then collected in a spin column, and processed as described above for AAVR PKD2 pull-downs.

### Retro-orbital injection and tissue preparation

Mice were retro-orbitally injected with AAVs (AAV9, PHP.eB, CAP-B10, eB.24, AAV9- B10) carrying CAG-GFP transgene. AAVs were administered under isoflurane anesthesia (1-3% in 95% O_2_ / 5% CO_2_, provided by nose cone at 1 L/min), followed by administration of 1-2 drops of 0.5% proparacaine to the corneal surface.

Three weeks post-injection, mice were euthanized via intraperitoneal injection of 100 mg/kg euthasol (pentobarbital sodium and phenytoin sodium solution, Virbac AH). Transcardial perfusion was performed using 30 mL of ice-cold heparinized 0.1M PBS pH 7.4, followed by an equal volume of ice-cold 4% paraformaldehyde (PFA) in 0.1M PBS. Collected organs were post-fixed in 4% PFA overnight at 4°C, rinsed, and preserved in 0.1M PBS with 0.05% sodium azide at 4 °C. Once tissues were fully equilibrated, they were embedded in O.C.T. Compound, flash-frozen using a dry ice-ethanol bath, and stored at -70 °C until sectioning. Tissues were prepared at 80 μm thickness using a cryostat (Leica Biosystems) and stored in 1x PBS at 4 °C until further processing.

### Immunohistochemistry

Immunohistochemistry was performed on free-floating tissue sections in 1x PBS. Sections were blocked with BlockAid Blocking Solution (ThermoFisher, B10710) containing 0.1% Triton X-100 (Sigma-Aldrich, #93443). This blocking buffer was also used for diluting primary and secondary antibodies. Tissue sections were incubated with primary antibodies overnight at 4 °C, followed by three 10-minute washes in 1x PBS. Subsequently, tissue sections were incubated with secondary antibodies for 2 hours at room temperature and washed three times for 10 min each with 1x PBS. For nuclear staining, tissue sections were incubated with 1/10000 Hoechst 33342 (ThermoFisher, H3570) in 1x PBS for 10 min, followed by three 10-min washes in PBS. For segmentation of neuronal cells, tissue sections were Nissl stained using 1/400 NeuroTrace 640/660 (ThermoFisher, N21483) in 1x PBS. After staining, tissues were washed twice for 1 hr each at room temperature, followed by one overnight wash at 4 °C in 1x PBS with 0.1% Triton X-100. After staining, sections were dried on slides, and Prolong Diamond Antifade Mountant (ThermoFisher, P36965) was applied to mount coverslips. The following antibodies were used in this study: rabbit anti-S100B (1/500, Abcam, ab52642), fluorophore-conjugated F(ab’)2 fragment secondary antibody (1/1000, Jackson ImmunoResearch, cat# 711-606-152).

### Tissue section imaging and analysis

For overview images of mouse brain and liver sections, a Keyence BZ-X710 epifluorescence microscope was used, with a 10x, 0.45 NA air objective.

For imaging of fields of view for quantification, a Zeiss LSM 880 with a 25x, 0.8 NA water immersion objective was used. Imaging settings were chosen to capture the full dynamic range of the signal without saturating pixels. When possible, laser power was adjusted before adjusting detector gain. Imaging settings were first optimized on control samples before imaging of experimental samples. Fields of view were chosen while imaging non-experimental channels (e.g. Hoechst or S100B). Three non-adjacent sections were imaged in the cortex, cerebellum, and liver to compensate for local heterogeneity in these tissues. For each section, three fields of view were taken to minimize heterogeneity within quantification.

For liver transduction quantification, cellular nuclei were segmented from the Hoechst channel in CellProfiler (v4.2.5; https://cellprofiler.org/)^67^. Segmented nuclei were then used as a mask on the GFP channel. The GFP intensity was quantified from the masked images. The images were then filtered into positive or negative cells, allowing for the percentage of GFP+ cells to be calculated.

Classification of cortical cells as neuronal GFP-positive or astrocytic GFP-positive was done using CellProfiler. Astrocytes were segmented using the S100B channel in CellProfiler, while neuronal cells were segmented from the Nissl channel using Cellpose (v3.0.7; https://www.cellpose.org/). Images were batch processed using napari (v0.4.19.post1; https://napari.org/stable/) and the serialcellpose plugin (v0.2.2; https://www.napari-hub.org/plugins/napari-serialcellpose). An Anaconda (v2.5.4; https://www.anaconda.com/) distribution of Python (v3.10.14; https://www.python.org/) was used. The GFP channel was segmented using the DAPI channel and then thresholded in CellProfiler. All thresholds were chosen based on distribution of mean GFP intensity in soma. In both S100B and Nissl samples the colocalization between GFP and the cell type marker was calculated in CellProfiler.

Statistical significance of all measured tropism was determined using a two-tailed unpaired t-test in GraphPad Prism(v10.3.1). For all statistical analysis, significance is represented as *p <0.05, **p < 0.01, ***p < 0.001, and ****p < 0.0001; non-significance, p > 0.05.

### Cell infectivity assay

Cell infectivity assays were performed as previously described^44,46^. Briefly, HEK293T cells (ATCC, CRL-3216) were maintained in Dulbecco’s Modified Eagle Medium (DMEM) supplemented with 5% fetal bovine serum (FBS), 1% nonessential amino acids (NEAA), and 100 U/mL penicillin-streptomycin, at 37 °C with humid air containing 5 % CO_2_. Cells were seeded in 6-well plates, and at 80% confluency, cells were transfected with 2.53 μg of plasmid DNA containing a LY6A expression sequence. Cells were then transferred to 96-well plates at 20% confluency in FluoroBrite™ DMEM supplemented with 0.5% FBS, 1% NEAA, 100 U/mL penicillin-streptomycin, 1× GlutaMAX, and 15 μM HEPES. Cells expressing LY6A were transduced with AAVs of interest at 1×10^9^ or 5×10^9^ vg per well in triplicate. 24 hours later, cells were imaged with a Keyence BZ-X700 microscope with a 4× objective. Images were quantified as previously described, using our custom Python-based image processing toolkit (https://github.com/GradinaruLab/in-vitro-transduction-assay.).

## Data availability statement

Cryo-EM maps and atomic models have been deposited with the accession codes EMDB:XXX and PDB:XXX (CAP-B10), EMDB:XXX and PDB:XXX (CAP-B10 - PKD2), EMDB:XXX and PDB:XXX (AAV9-X1), EMDB:XX and PDB:XXX (AAV9-X1.1), EMDB:XXX and PDB:XXX (AAV9-X1.1 - PKD2), EMDB:XXX and PDB:XXX (AAV9-B10), and EMDB:XXX and PDB:XXX (eB.24).

## Acknowledgements

We thank all Gradinaru lab and CLOVER staff members for their mindful help. We thank Alex Jin Chung, and Yaping Lei for technical support. We thank Catherine Oikonomou for help with manuscript editing. All cryo-EM data were collected at the Caltech cryo-EM facility. Biolayer interferometry data were collected in the Caltech Protein Expression Center (PEC). We are especially thankful to Songye Chen (Caltech cryo-EM facility director) and Jost Vielmetter (Caltech PEC director) for supporting our data collection. We thank Travis Bainbridge and Beyza Bulutoglu for helpful discussions about optimizing BLI for AAVs. This project was supported by the Beckman Institute CLOVER Center (to T.F.S. and V.G.), NIH PIONEER DP1NS111369 (to V.G.), and NIH BRAIN Initiative Armamentarium UF1MH128336 (to V.G. and T.F.S.).

## Contributions to authorship

S.J., and V.G. conceived and T.B., S.J., T.S, and V.G designed the study. T.B. performed cryo-EM data collection and processing. T.B. and S.J. analyzed structure data. S.J. performed and analyzed BLI experiments. T.B., S.J., and N.A. performed molecular cloning. S.J. performed receptor protein production. C.C. performed size exclusion chromatography. S.J. performed pull-down assays. I.G. performed cell infectivity assays. G.C., B.B., and F.R. performed mouse injections, perfusions, and tissue staining. G.C. and S.J. performed Keyence imaging. T.B. performed confocal imaging and quantified liver de-targeting and brain tropism. T.B. and S.J. wrote the manuscript and prepared the figures. V.G. supervised all aspects of the work.

## Supplementary Figure legends

**Supplementary Figure 1:**

**PHP.eB and CAP-B10 interaction with PKD2. (a)** PHP.eB has a 7-mer insertion at VR-VIII (^1’^TLAVPFK^7’^) between AA 588 and 589, and two adjacent point mutations (^587^AQ^588^ to ^587^DG^588^). In a previous study, we identified that the acidic residue D587 forms a hydrophilic interaction with the basic residue K7’ from the 7-mer, which we termed the ‘lysine-lever’. This interaction creates structural tension, causing VR-VIII to bend inward. The inwardly bent conformation introduces a steric clash with AAVR-PKD2, thereby reducing the binding affinity. AAV9 (PDB: 3UX1^19^) is shown in purple. PHP.eB (PDB: 7UD4^21^) is shown in blue. In PHP.eB - PKD2 complex (PDB: 7WQP^38^), PHP.eB is shown in blue, and PKD2 is presented with electrostatic mapping. **(b)** Illustration of how VR-IV and VR-VIII from a single AAV monomer interact with two PKD2 molecules. The three AAV monomers which compose the 3-fold face are shown in white, light gray, and dark gray. AAVR-PKD2 is red. VR-IV and VR-VIII are blue and green, respectively. VR-IV and VR-VIII are positioned between two neighboring monomers, allowing them to interact with both. The posterior PKD2 interacts with the protruding end of VR-VIII, including the PHP.eB 7-mer, while the anterior PKD2 interacts with VR-IV and VR-VIII.

**Supplementary Figure 2:**

**Resolution of all AAV structures determined in this study. (a)** Gold standard FSC curves with indicated resolution at 0.143 cutoff (left), azimuth elevation plots (middle), and local resolution of each cryo-EM map (right). **(b, c)** Electron density maps of VR-IV (**b**) and VR-VIII (**c**) regions of AAV structures with associated atomic models.

**Supplementary Figure 3:**

**Cryo-EM single particle reconstruction processing pipeline.** All cryo-EM structures in this study were processed using cryoSPARC (v4.1). Details of steps and parameters used are described in the Methods and Supplementary Table 1. Briefly, raw movies were binned by 2 and gain and motion corrected in CryoSPARC(v4.1)^63^ CTF estimation was performed in CryoSPARC . Initial particle picks were generated with blob picker and were inspected to eliminate bad picks. 2D classification with 100 classes was then used to exclude remaining junk particles. Multiple rounds of refinement jobs in I1 symmetry were performed on good particles to construct the final map for model building. A model of the VP1 capsid protein was built based on AAV9 as a template (PDB: 3UX1^19^), using Phenix and COOT^39,40,64^.

**Supplementary Figure 4:**

**Supporting structure information for Figure 2. (a, b)** Possible hydrogen bonding interactions between PKD2 and PHP.eB VR-IV (PDB: 7WQP). **(a)** Rotamer screening identified the most favorable rotamer position of Q456 (prevalence score: 0.079, Rank 1), which can form a hydrogen bond with E418 of PKD2 at a distance of 3.07 Å. Additionally, this rotamer can establish a hydrogen bond with the carbonyl oxygen of the peptide bond in E418. **(b)** Rotamer screening also identified a less favorable rotamer position of Q456 (prevalence score: 0.007, Rank 29), which can form a hydrogen bond with N496 at a distance of 3.53 Å. This greater distance compared to the hydrogen bonding with E418 suggests a weaker or less stable interaction. **(c–e)** Cryo-EM structure of our designed AAV, eB.24 **(c)**, and a cryo-EM-based atomic model of eB.24 overlaid with PHP.eB (PDB: 7UD4) and CAP-B10 (this study) **(d, e)**. The backbone structures of VR-IV are identical in eB.24 and PHP.eB, whereas CAP-B10 differs, indicating that the S454A and Q456T mutations are insufficient to alter the overall loop structure. PHP.eB, eB.24, and CAP-B10 all share an identical backbone structure of VR-VIII.

**Supplementary Figure 5:**

**AAV-receptor pull-down assay. (a)** Schematic of pull-down assay, using receptor as bait and AAV as prey. For protein receptors, receptors tagged with 6xHis were immobilized on agarose beads, and the amount of AAV captured by the receptor wa analyzed using Western blot. For D-galactose receptor, D-galactose pre-loaded agarose beads were used. To prevent signal saturation, either ½ or ⅓ of the input was loaded. The amount of AAV pulled down was normalized to the input and is represented below each lane**. (b)** Pull-down of AAV9, PHP.eB, and CAP-B10 by PKD2. CAP-B10 was pulled down the least, consistent with the relative ranking of binding affinities measured by BLI. **(c-e)** Pull-down of PHP.eB, eB.24, and CAP-B10 by PKD2, LY6A, or D-galactose. **(c)** The degree of VR-IV modification of the capsids showed an inverse correlation with PKD2 binding strength. **(d,e)** LY6A and D-galactose binding did not show a similar trend, suggesting that VR-IV modifications specifically impact AAVR-PKD2 interaction.

**Supplementary Figure 6:**

**Supporting structure information for Figures 3 & 4. (a)** Cryo-EM map of AAV9-X1. **(b,c)** Atomic model of AAV9-X1 (brown), showing whole monomer, VR-IV, and VR-VIII regions, superimposed on CAP-B10 (orange), AAV9-X1.1 (yellow), and PKD2-bound AAV9-X1.1 (blue) to highlight structural differences and similarities. AAV9-X1 exhibits a distinct VR-IV structure compared to CAP-B10 and AAV9-X1.1. The VR-IV loop structure is similar between CAP-B10 and AAV9-X1.1, and is deformed when PKD2 binds. **(d)** Cryo-EM reconstruction of AAV9-B10 structure. **(e,f)** Atomic model of AAV9-B10, showing the full capsid, VR-IV, and VR-VIII regions, built based on the cryo-EM structure (blue). These models are superimposed with CAP-B10 (orange) and AAV9 (gray) to illustrate structural differences and similarities. AAV9-B10 and CAP-B10 share an identical VR-IV loop structure.

**Supplementary Figure 7:**

**Cell infectivity assays (a)** Cell infectivity assay representing the transduction efficiency of each capsid in HEK293T cells either untransfected or transfected with a construct to express LY6A on their membranes. AAV9, AAV9-B10, PHP.eB, eB.24, and CAP-B10 were tested at a dosage of 1×10^9^ vg / well. Color indicates extent of infection and size indicates total brightness per signal area. Data were quantified from triplicate experiments. **(b)** Rescaled comparison of AAV9 and AAV9-B10 to highlight differences in transduction efficiency. AAV9-B10 showed lower potency than AAV9. **(c)** Rescaled comparison of PHP.eB, eB.24, and CAP-B10 in untransfected HEK293T cells (lacking LY6A), revealing a weak inverse correlation between PKD2 binding affinity and transduction efficiency, though basal transduction levels were low. **(d)** Cell infectivity of PHP.eB, eB.24, and CAP-B10 at a higher dosage of 5×10^9^ vg per well. The inverse correlation between PKD2 binding affinity and cell transduction potency became more apparent. However, this trend was not observed in LY6A-expressing cells, suggesting that alternative receptors can override the effects of reduced PKD2 binding. **(e)** Representative fluorescence images of HEK293T cells expressing eGFP delivered by each capsid.

