## Supplementary material for "Structural basis of liver de-targeting and neuronal tropism of CNS-targeted AAV capsids": Table1

| Capsid | AAVR-PKD2 affinity rank | K_D_ against AAVR-PKD2 (μM) | Systemic Liver transduction | Mouse CNS tropism | Engineered Receptor | Receptor Distribution | | Ref. |
| --- | --- | --- | --- | --- | --- | --- | --- | --- |
|  |  |  |  |  |  | Liver | CNS |  |
| AAV9-X1 | 1 | 5.87 | +++ | Endothelial | LRP6 | O | O | 13. |
| AAV9  (wild-type) | 2 | 11.40 | +++ | Weak | - |  |  | 4. |
| PHP.eB | 3 | 14.22 | ++ | Neuronal & Astrocytic | LY6A | X | O | 11. |
| AAV9-X1.1 | 4 | 18.85 | +++ | Endothelial | LRP6 | O | O | 13. |
| eB.24 | 5 | 24.15 | + | Neuronal &  less Astrocytic | LY6A | X | O | This study |
| AAV9-B10 | 6 | 46.90 | ++ | Weak | - |  |  | This study |
| CAP-B10 | 7 | 60.63 | Very low | Highly Neuronal | LY6A | X | O | 12. |

Table 1: AAVR-PKD2 binding affinity ranking, and *in vivo* tropism of capsids used in this study.
