## Supplementary Figures for "Structural basis of liver de-targeting and neuronal tropism of CNS-targeted AAV capsids"

### Supplementary Figure 1.

(a)

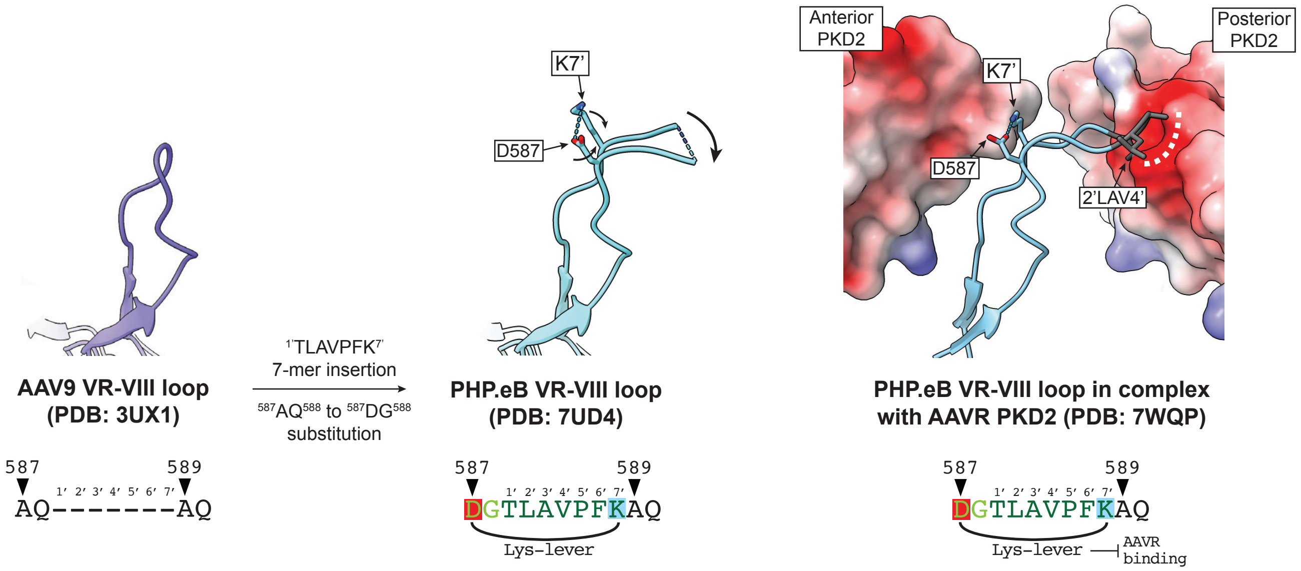


(b)

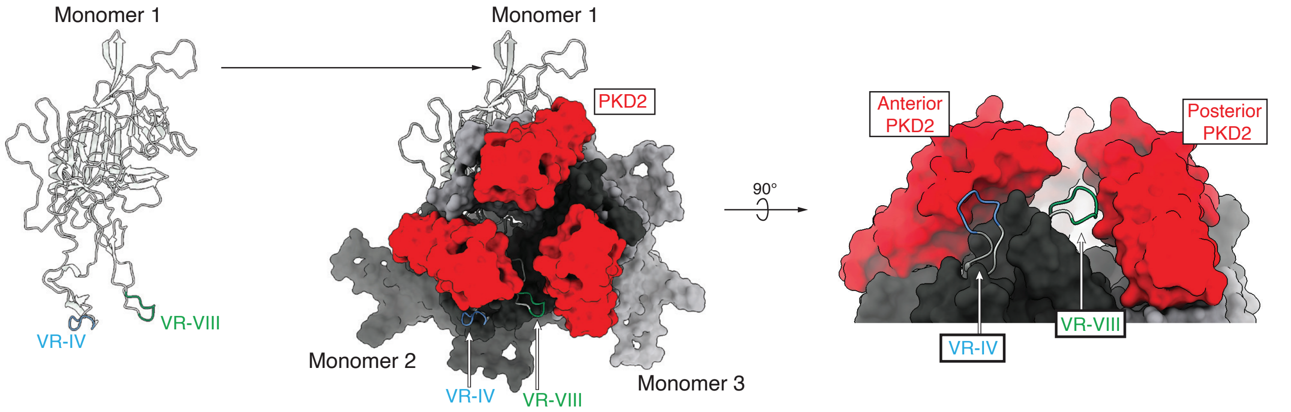

### Supplementary Figure 2.

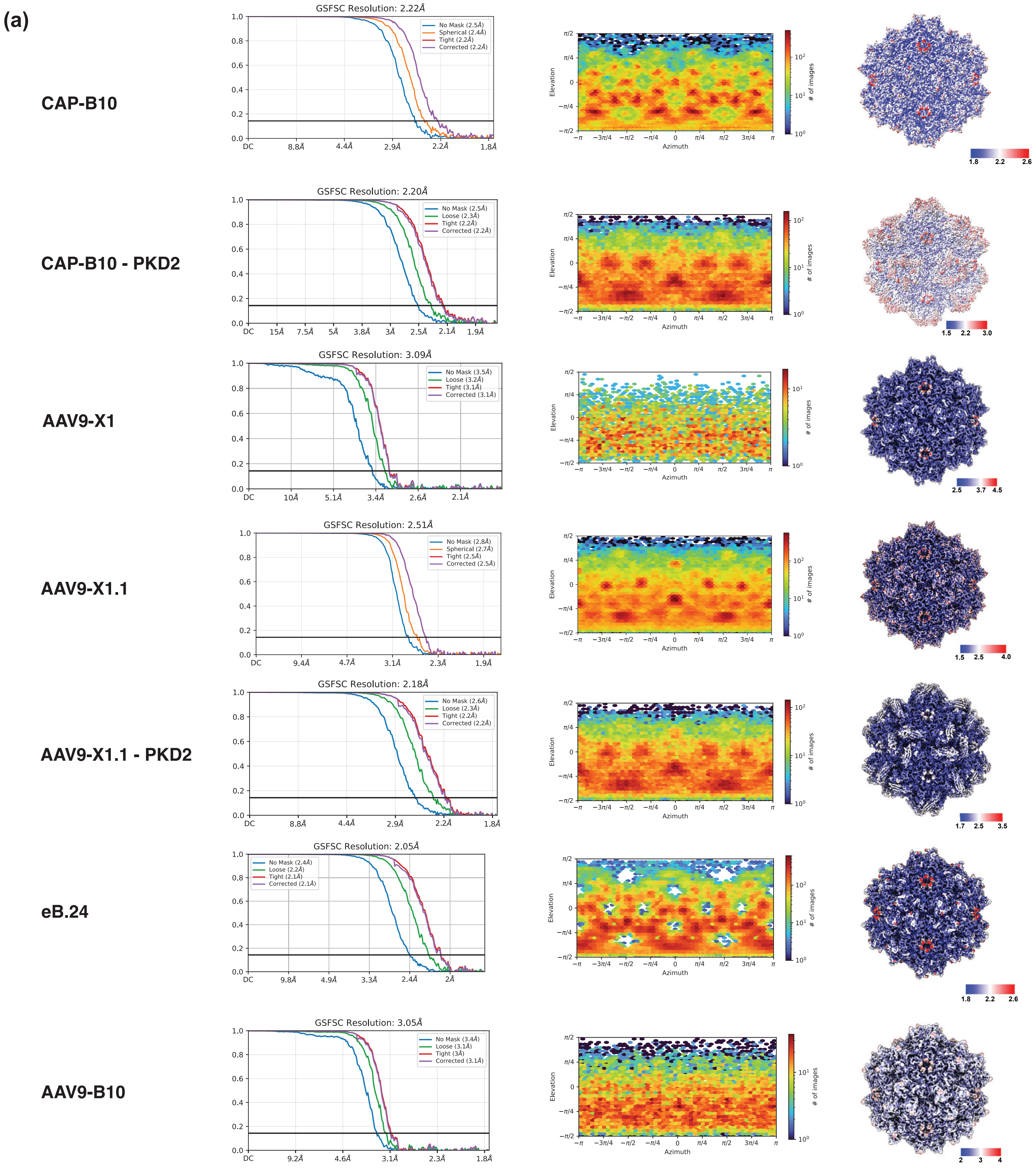

#### Supplementary Figure 2.

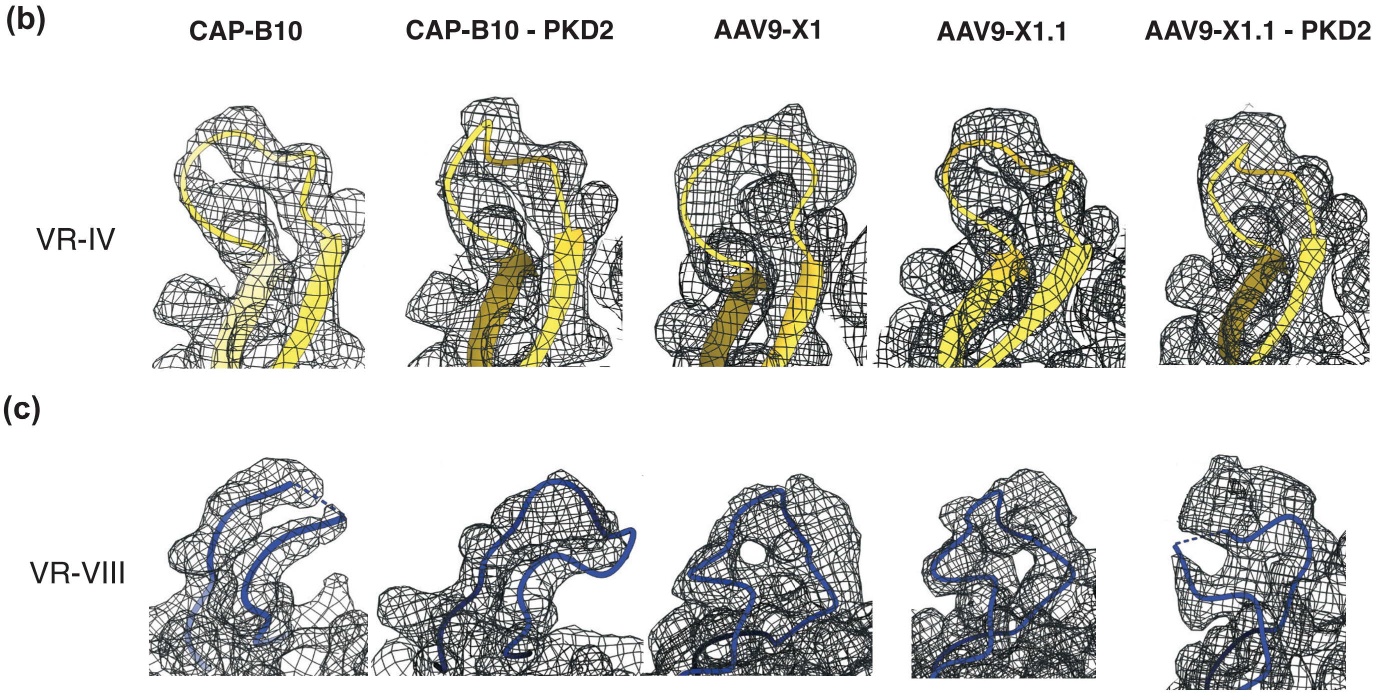

#### Supplementary Figure 3.

(a)

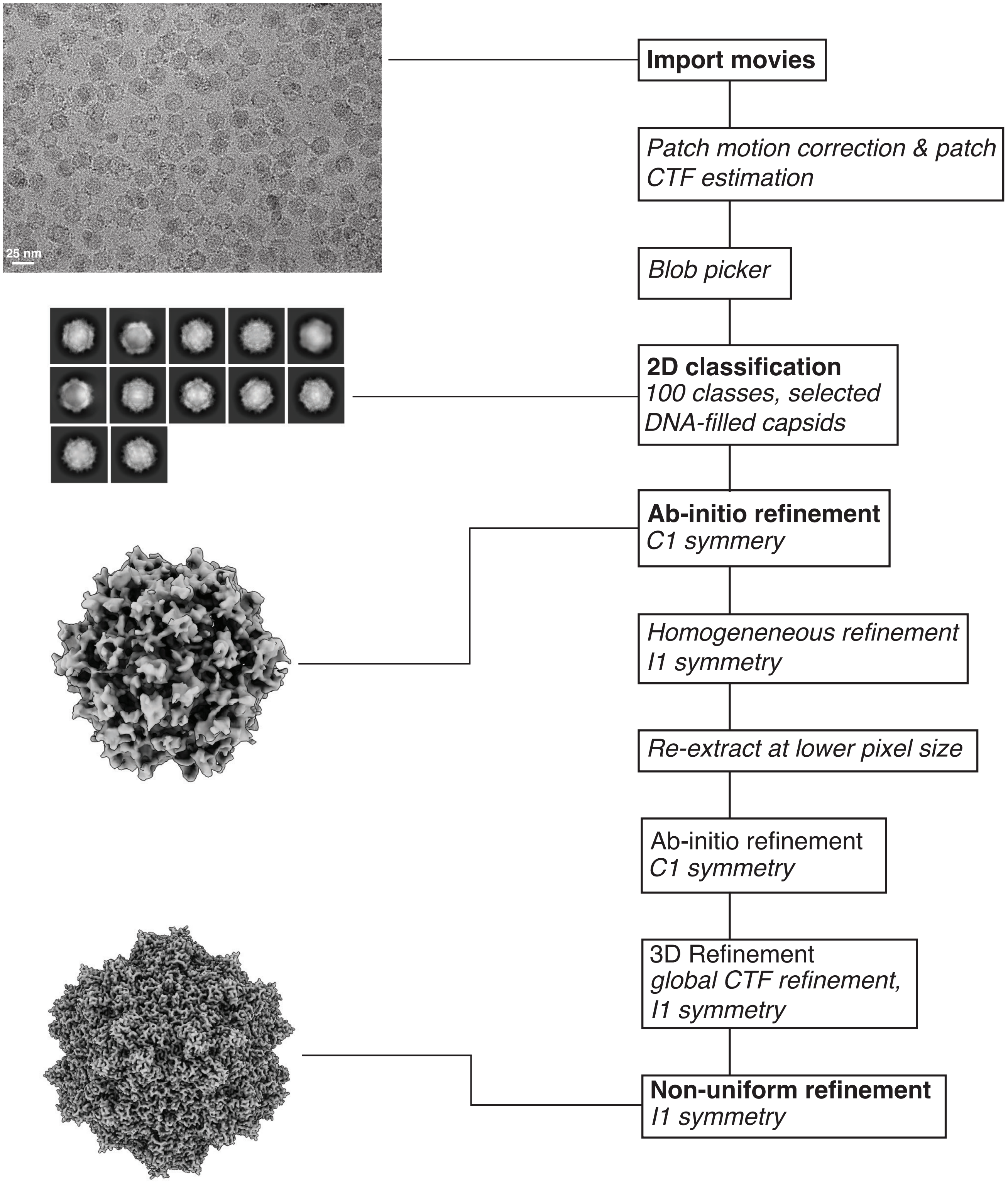

### Supplementary Figure 4.

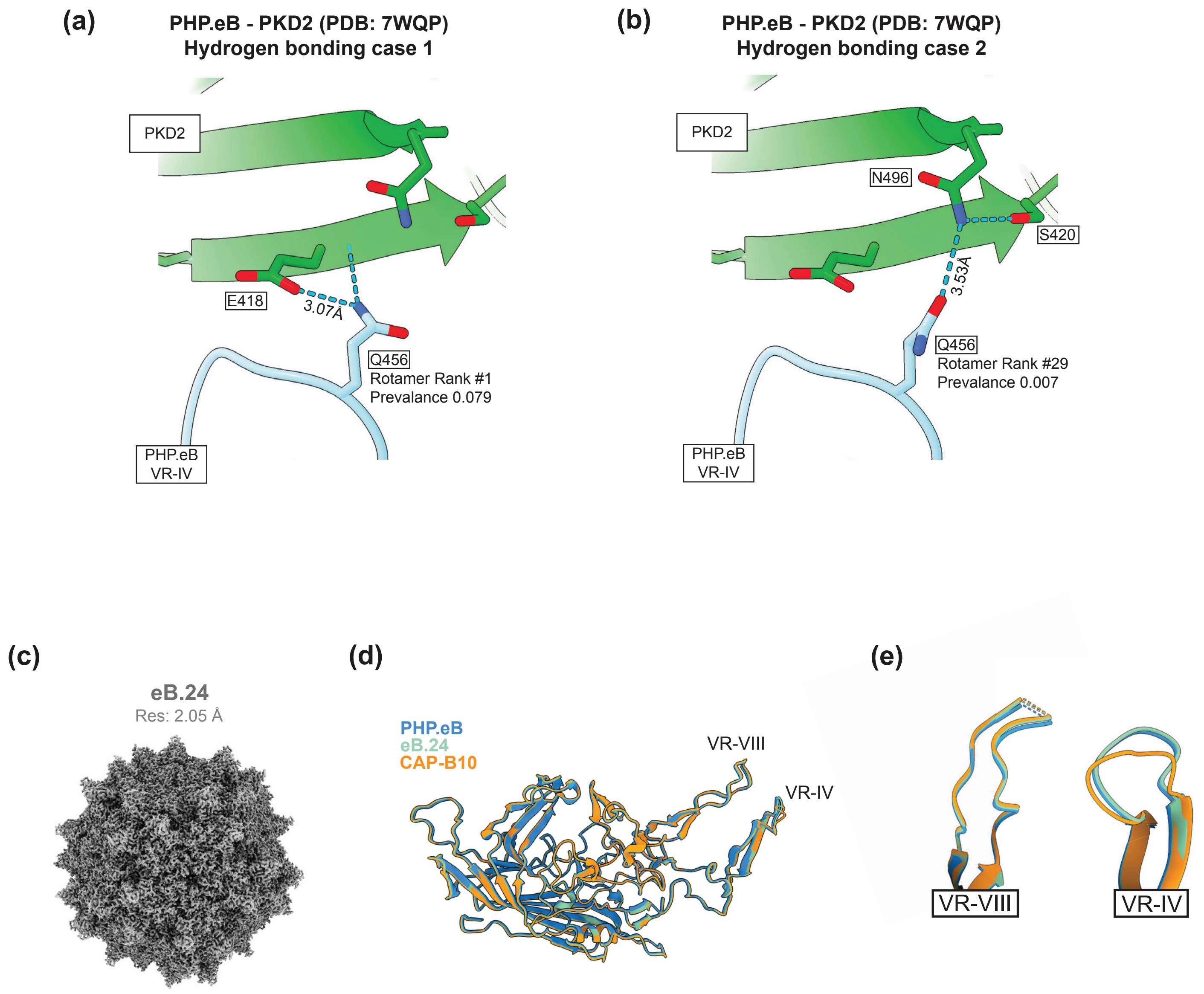

### Supplementary Figure 5.

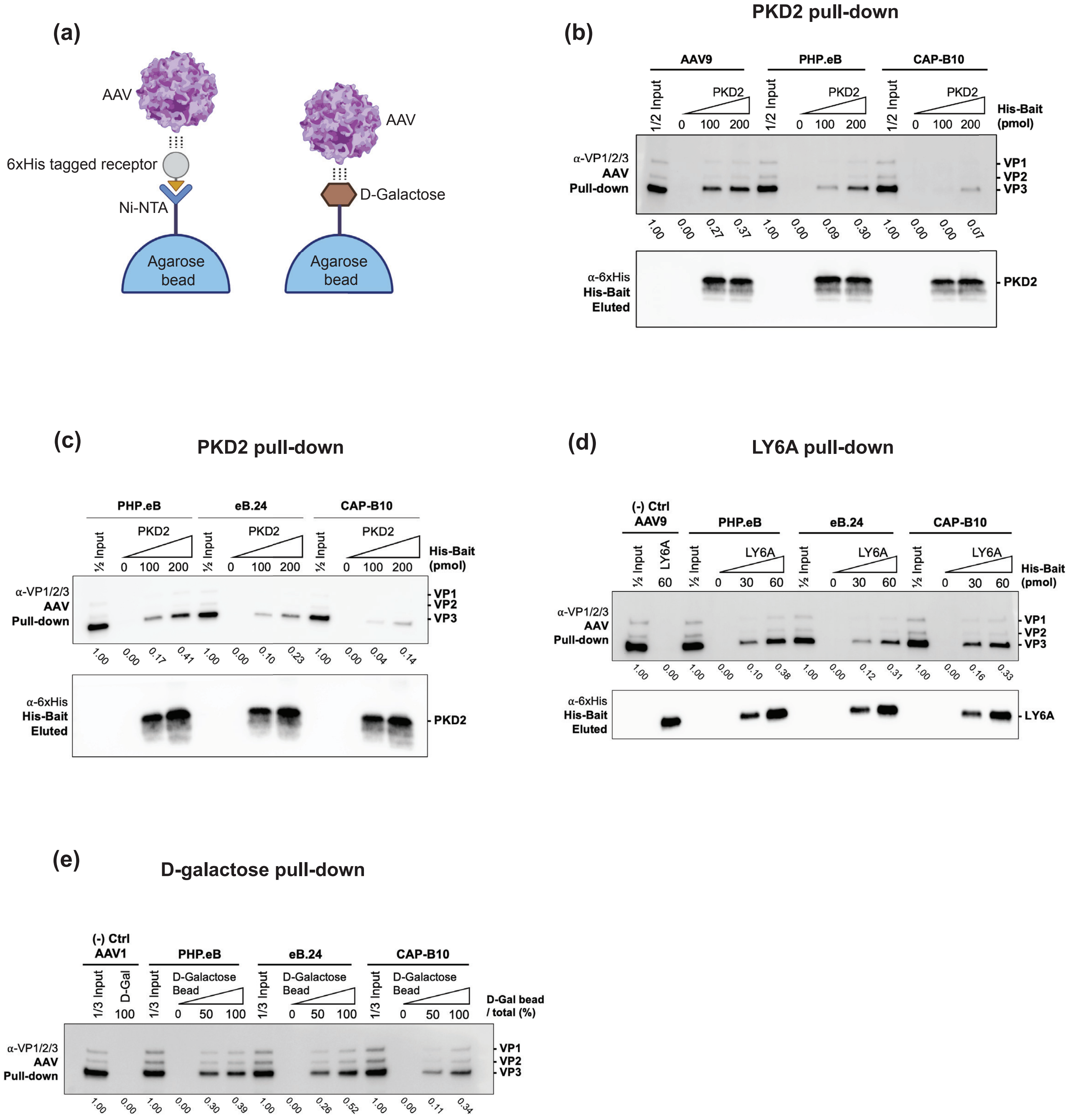

**Supplementary Figure 6.**

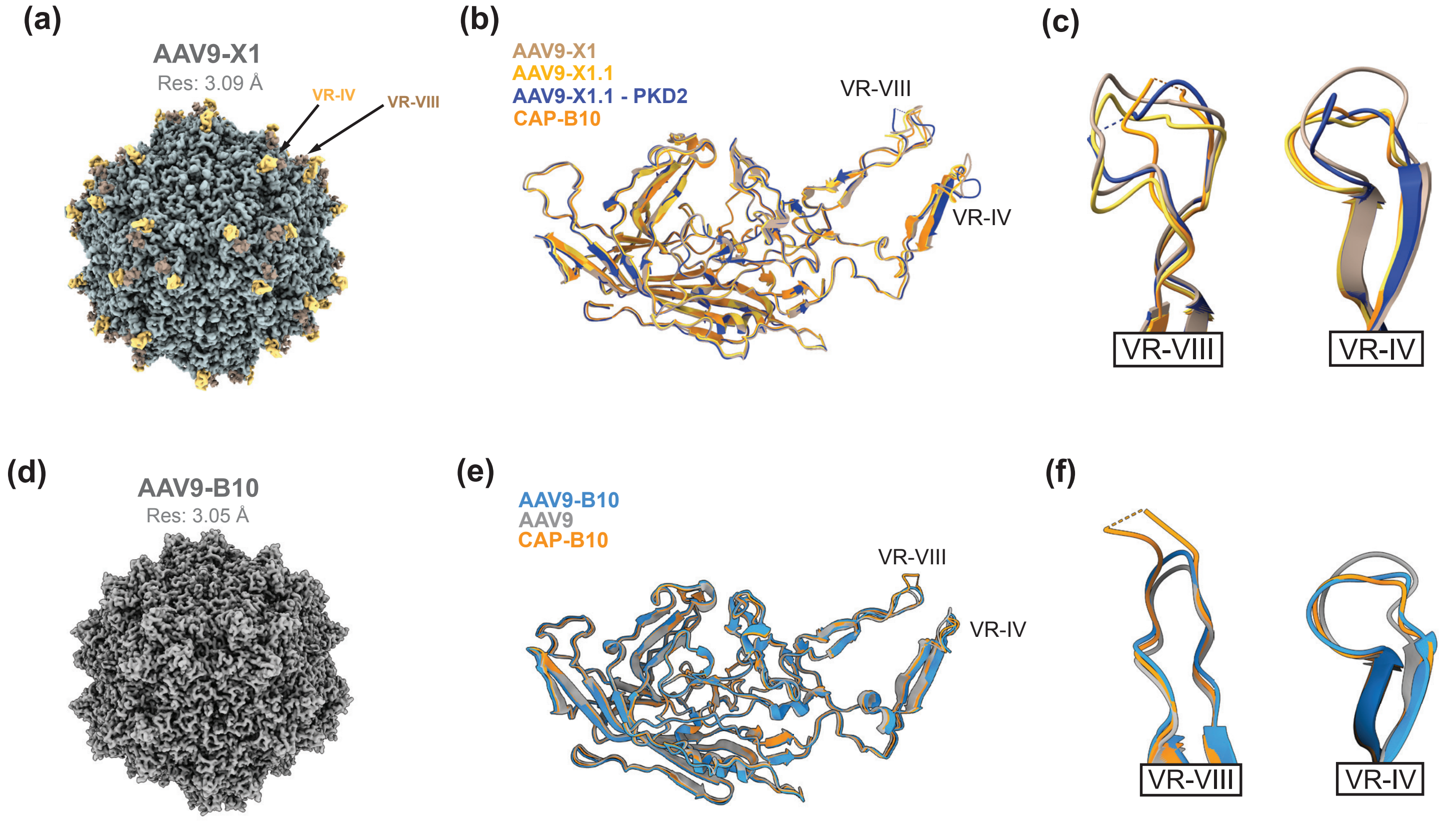

### Supplementary Figure 7.

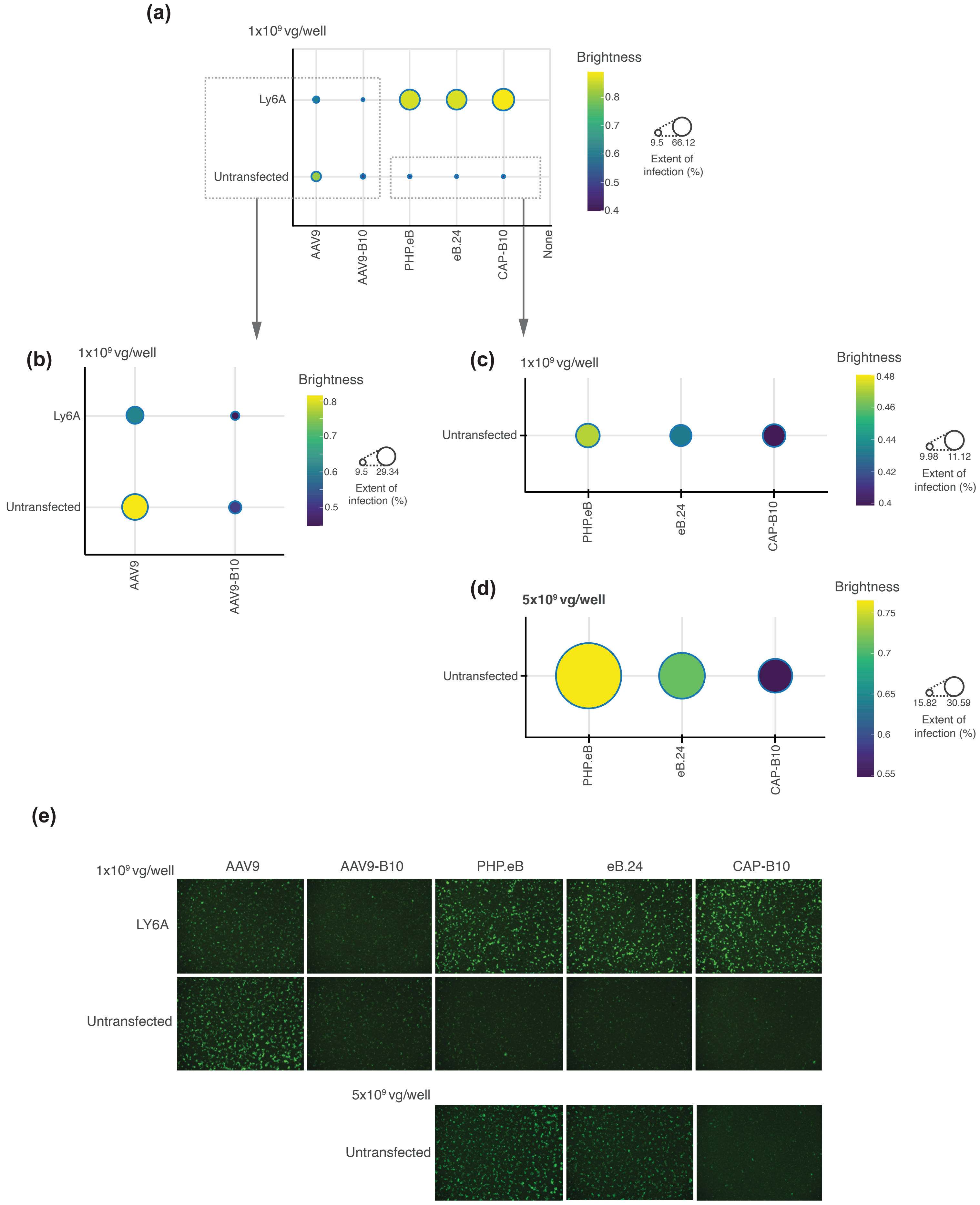
