## Supplementary Tables for "Structural basis of liver de-targeting and neuronal tropism of CNS-targeted AAV capsids"

| CAP-B10  EMDB-X  PDB X | |
| --- | --- |
| Data collection and processing: | |
| Magnification | x105,000 |
| Voltage(kV) | 300 |
| Electron exposure(e^-^/Å^2^) | 60 |
| Defocus range(µm) | -1 to -3 |
| Pixel size (Å) | 0.42 |
| Symmetry imposed | I1 |
| Initial particle images (no.) | 386,730 |
| Final particle images (no.) | 104,127 |
| Map resolution (Å)  FSC threshold: 0.143 | 2.22 |
| Refinement: | |
| Initial model used | 7UD4 |
| Correlation coefficient (CC_mask_) | 0.95 |
| R.M.S devations | |
| Bond lengths (Å) | 0.003(0) |
| Bond angles (˚) | 0.501(0) |
| Validation | |
| MolProbity score | 1.03 |
| Clashscore | 1.24 |
| Poor rotamers (%) | 0.00 |
| Ramachandran plot | |
| Favored (%) | 96.91 |
| Allowed (%) | 3.09 |
| Disallowed (%) | 0.00 |

| CAP-B10 PKD2 complex  EMDB-X  PDB X | |
| --- | --- |
| Data collection and processing: | |
| Magnification | x105,000 |
| Voltage(kV) | 300 |
| Electron exposure(e^-^/Å^2^) | 60 |
| Defocus range(µm) | -1 to -3 |
| Pixel size (Å) | 0.42 |
| Symmetry imposed | I1 |
| Initial particle images (no.) | 354,088 |
| Final particle images (no.) | 56,972 |
| Map resolution (Å)  FSC threshold: 0.143 | 2.20 |
| Refinement: | |
| Initial model used | 7WQP |
| Correlation coefficient (CC_mask_) | 0.91 |
| R.M.S devations | |
| Bond lengths (Å) | 0.003(0) |
| Bond angles (˚) | 0.532(0) |
| Validation | |
| MolProbity score | 1.45 |
| Clashscore | 2.94 |
| Poor rotamers (%) | 0.74 |
| Ramachandran plot | |
| Favored (%) | 94.62 |
| Allowed (%) | 5.38 |
| Disallowed (%) | 0 |

| AAV9-X1  EMDB-X  PDB X | |
| --- | --- |
| Data collection and processing: | |
| Magnification | x105,000 |
| Voltage(kV) | 300 |
| Electron exposure(e^-^/Å^2^) | 60 |
| Defocus range(µm) | -1.5 to -3 |
| Pixel size (Å) | 0.42 |
| Symmetry imposed | I1 |
| Initial particle images (no.) | 835,394 |
| Final particle images (no.) | 11,999 |
| Map resolution (Å)  FSC threshold: 0.143 | 3.09 |
| Refinement: | |
| Initial model used | 7UD4 |
| Correlation coefficient (CC_mask_) | 0.92 |
| R.M.S devations | |
| Bond lengths (Å) | 0.007(13) |
| Bond angles (˚) | 1.114(34) |
| Validation | |
| MolProbity score | 1.02 |
| Clashscore | 1.11 |
| Poor rotamers (%) | 0.22 |
| Ramachandran plot | |
| Favored (%) | 96.75 |
| Allowed (%) | 3.25 |
| Disallowed (%) | 0 |
| AAV9-X1.1  EMDB-X  PDB X | |
| Data collection and processing: | |
| Magnification | x105,000 |
| Voltage(kV) | 300 |
| Electron exposure(e^-^/Å^2^) | 60 |
| Defocus range(µm) | -1.5 to -3 |
| Pixel size (Å) | 0.42 |
| Symmetry imposed | I1 |
| Initial particle images (no.) | 449,282 |
| Final particle images (no.) | 112,046 |
| Map resolution (Å)  FSC threshold: 0.143 | 2.51 |
| Refinement: | |
| Initial model used | 7UD4 |
| Correlation coefficient (CC_mask_) | 0.93 |
| R.M.S devations | |
| Bond lengths (Å) | 0.002 (0) |
| Bond angles (˚) | 0.459 (0) |
| Validation | |
| MolProbity score | 0.73 |
| Clashscore | 0.12 |
| Poor rotamers (%) | 0 |
| Ramachandran plot | |
| Favored (%) | 96.94 |
| Allowed (%) | 3.06 |
| Disallowed (%) | 0.00 |

| AAV9-X1.1 PKD2 complex  EMDB-X  PDB X | |
| --- | --- |
| Data collection and processing: | |
| Magnification | x105,000 |
| Voltage(kV) | 300 |
| Electron exposure(e^-^/Å^2^) | 60 |
| Defocus range(µm) | -1.5 to -3 |
| Pixel size (Å) | 0.42 |
| Symmetry imposed | I1 |
| Initial particle images (no.) | 734,883 |
| Final particle images (no.) | 81,365 |
| Map resolution (Å)  FSC threshold: 0.143 | 2.18 |
| Refinement: | |
| Initial model used | 7WQP |
| Correlation coefficient (CC_mask_) | 0.93 |
| R.M.S devations | |
| Bond lengths (Å) | 0.003 (0) |
| Bond angles (˚) | 0.515(1) |
| Validation | |
| MolProbity score | 1.54 |
| Clashscore | 2.41 |
| Poor rotamers (%) | 2.4 |
| Ramachandran plot | |
| Favored (%) | 96.56 |
| Allowed (%) | 3.44 |
| Disallowed (%) | 0 |

| eB.24  EMDB-X  PDB X | |
| --- | --- |
| Data collection and processing: | |
| Magnification | x105,000 |
| Voltage(kV) | 300 |
| Electron exposure(e^-^/Å^2^) | 60 |
| Defocus range(µm) | -1.5 to -3 |
| Pixel size (Å) | 0.42 |
| Symmetry imposed | I1 |
| Initial particle images (no.) | 433,853 |
| Final particle images (no.) | 238,315 |
| Map resolution (Å)  FSC threshold: 0.143 | 2.05 |
| Refinement: | |
| Initial model used | 7UD4 |
| Correlation coefficient (CC_mask_) | 0.91 |
| R.M.S devations | |
| Bond lengths (Å) | 0.002 (0) |
| Bond angles (˚) | 0.398 (0) |
| Validation | |
| MolProbity score | 1.06 |
| Clashscore | 2.48 |
| Poor rotamers (%) | 1.10 |
| Ramachandran plot | |
| Favored (%) | 98.26 |
| Allowed (%) | 1.74 |
| Disallowed (%) | 0 |
| AAV9-B10  EMDB-X  PDB X | |
| Data collection and processing: | |
| Magnification | X45,000 |
| Voltage(kV) | 200 |
| Electron exposure(e^-^/Å^2^) | 60 |
| Defocus range(µm) | -1.5 to -3 |
| Pixel size (Å) | 0.90 |
| Symmetry imposed | I1 |
| Initial particle images (no.) | 112,168 |
| Final particle images (no.) | 26,411 |
| Map resolution (Å)  FSC threshold: 0.143 | 3.05 |
| Refinement: | |
| Initial model used | 7UD4 |
| Correlation coefficient (CC_mask_) | 0.92 |
| R.M.S devations | |
| Bond lengths (Å) | 0.005 (0) |
| Bond angles (˚) | 0.545 (0) |
| Validation | |
| MolProbity score | 1.21 |
| Clashscore | 2.24 |
| Poor rotamers (%) | 0.00 |
| Ramachandran plot | |
| Favored (%) | 96.71 |
| Allowed (%) | 3.29 |
| Disallowed (%) | 0 |

Supplementary Table 2. Average binding kinetic constants of AAVs and AAVR-PKD2.

|  | PKD2 binding affinity rank | K_a_ (1/Ms) | K_d_ (1/s) |
| --- | --- | --- | --- |
| X1 | 1 | 1.27± 0.56 E4 | 0.73± 0.12 E-1 |
| AAV9 | 2 | 1.17± 0.16 E4 | 1.32± 0.16 E-1 |
| eB | 3 | 0.86± 0.19 E4 | 1.19± 0.24 E-1 |
| X11 | 4 | 0.86± 0.2 E4 | 1.57± 0.29 E-1 |
| eB.24 | 5 | 0.7± 0.12 E4 | 1.69± 0.36 E-1 |
| AAV9B10 | 6 | 0.52± 0.04 E4 | 2.44± 0.08 E-1 |
| B10 | 7 | 0.55± 0.03 E4 | 3.35± 0.2 E-1 |
